# Endothelial MICU1 protects against vascular inflammation and atherosclerosis by inhibiting mitochondrial calcium uptake

**DOI:** 10.1101/2024.08.22.609192

**Authors:** Lu Sun, Ruixue Leng, Qingze He, Meiming Su, Zhidan Zhang, Zhenghong Liu, Zhihua Wang, Hui Jiang, Monan Liu, Li Wang, Yuqing Huo, Clint L Miller, Maciej Banach, Yu Huang, Paul C Evans, Jaroslav Pelisek, Giovanni G Camici, Bradford C Berk, Stefan Offermanns, Junbo Ge, Suowen Xu, Jianping Weng

**Affiliations:** Department of Endocrinology, Institute of Endocrine and Metabolic Disease, The First Affiliated Hospital of USTC, Division of Life Sciences and Medicine, Clinical Research Hospital of Chinese Academy of Sciences (Hefei), University of Science and Technology of China, Hefei, Anhui, 230001, China; Department of Pharmacy, The First Affiliated Hospital of USTC, Division of Life Sciences and Medicine, University of Science and Technology of China, Hefei, Anhui, 230001, China; Department of Epidemiology and Biostatistics, School of Public Health, Anhui Medical University,Hefei, China; Department of Biomedical Sciences, City University of Hong Kong, China; Vascular Biology Center, Medical College of Georgia, Augusta University, Augusta, Georgia, 30912, USA; Center for Public Health Genomics, University of Virginia, Charlottesville, VA, USA; Department of Preventive Cardiology and Lipidology, Medical University of Lodz (MUL), Rzgowska 281/289, 93-338, Lodz, Poland; Centre for Biochemical Pharmacology, William Harvey Research Institute, Barts and The London Faculty of Medicine and Dentistry, Queen Mary University of London, London, UK; Department of Vascular Surgery, University Hospital Zurich, Zurich, Switzerland; Center for Molecular Cardiology, Schlieren Campus, University of Zurich, Wagistrasse 12, 8952 Schlieren, Switzerland; Aab Cardiovascular Research Institute, Department of Medicine, University of Rochester School of Medicine and Dentistry, Rochester, NY, USA; Max Planck Institute for Heart and Lung Research, Department of Pharmacology, Ludwigstr. 43, 61231, Bad Nauheim, Germany; Department of Cardiology, The First Affiliated Hospital of USTC, Division of Life Sciences and Medicine, University of Science and Technology of China, Hefei, Anhui, 230001, China

**Keywords:** atherosclerosis, calcium, endothelial inflammation, MICU1, oxidative stress

## Abstract

Atherosclerosis is triggered by endothelial activation and vascular inflammation, which is closely related to mitochondrial dysfunction. Mitochondrial calcium uptake 1 (MICU1), as the gatekeeper of mitochondrial Ca^2+^ homeostasis, is a critical player in mitochondrial function and implicated in a plethora of pathophysiological conditions. However, the role of MICU1 in the pathogenesis of vascular inflammation and atherosclerosis is unknown. We ask whether endothelial MICU1 can prevent vascular inflammation and atherosclerosis by maintaining mitochondrial homeostasis. We observed that vascular inflammation in response to LPS was aggravated in EC-specific *Micu1* knockout mice (*Micu1*^ECKO^) and reduced in EC-specific *Micu1* transgenic mice (*Micu1*^ECTg^). Furthermore, hypercholesterolemic *Micu1*^ECKO^ mice also showed accelerated formation of atherosclerotic plaques, while *Micu1*^ECTg^ mice protected against atherosclerosis. Mechanistically, MICU1 regulated mitochondrial Ca^2+^ influx, thereby reducing the expression of the mitochondrial deacetylase SIRT3 and the ensuing deacetylation of SOD2, leading to the burst of mitochondrial reactive oxygen species (mROS). Of clinical relevance, we identified decreased MICU1 level in endothelial layer of human atherosclerotic plaques, as well as in primary human aortic endothelial cells (HAECs) exposed to serum from patients with coronary artery diseases. Two-sample Wald ratio Mendelian randomization further revealed that increased expression of MICU1 was associated with decreased risk of coronary artery bypass grafting (CABG). Our findings unravel a critical and unrecognized role of MICU1 in preventing vascular inflammation and atherosclerosis by maintaining mitochondrial Ca^2+^ homeostasis.

## Introduction

Atherosclerosis, a chronic vascular inflammatory disease, is the leading cause of coronary artery disease (CAD) (1). Vascular inflammation is an early independent risk factor for atherosclerosis and CAD, and targeting inflammation has been reported to be next frontier for atherosclerosis treatment (2, 3). Antiatherosclerosis have shifted focus beyond lipid-lowering strategies towards addressing residual inflammatory risks involved in major adverse cardiovascular events (4). Endothelium is the primary target for vascular inflammation. Upon activation, it responds through an increased expression of leukocyte adhesion molecules and pro-inflammatory cytokines, which consequently leads to chronic inflammation and promotes atherosclerosis (5). Thus, the protection of endothelium from inflammation is a fundamental strategy in preventing vascular inflammatory diseases and atherosclerosis.

Mitochondria play an important role in cellular metabolism and energy generation in mammalian physiology. Finetuning of mitochondria Ca^2+^ level is central to orchestrating mitochondrial homeostasis (6). Mitochondria are a principal source of ATP but also reactive oxygen species (ROS), which are upstream of multiple kinases and cytokines. Overload of Ca^2+^ in mitochondria leads to excessive accumulation of mitochondrial ROS (mROS), accompanied by mitochondria dysfunction, energy metabolic disorder, cell damage and even death (6, 7). Besides, mROS is closely associated with inflammation (8, 9), which underscores the potential to suppress inflammatory response by controlling mROS.

MICU1 is the first reported member of the mitochondria calcium uniporter (MCU) complex (10). The MCU complex consists of mitochondria calcium uptake (MCU) as Ca^2+^ uptake channel (11, 12), MICU1-3 as regulatory subunits (10, 13), MCUb and essential MCU regulator (EMRE) (14, 15). Mitochondrial Ca^2+^ accumulation is regulated by MCU complex, as MCU oligomerizes to form a channel structure and MICU1 senses Ca^2+^ via its two EF-hands (10, 16). Increased expression and formation of MCU result in mitochondrial Ca^2+^ overload, inducing cell death and activation of NLRP3 inflammasome pathway (17–19). Consequently, MCU depletion might restore dynamin-related protein 1-mediated mitochondrial fission and thus facilitates the clearance of apoptotic cells involved in atherosclerosis (20). MICU1 serves as an essential gatekeeper of MCU in a closed conformation at low cytosolic Ca^2+^ concentrations to regulate mitochondria signaling and function (10, 16, 21–23). Recent evidence has revealed that MICU1 is also essential for adaptation of liver regeneration, lung alveolar type 2 cell plasticity and against neurodegeneration (24–26). However, the specific role of MICU1 in endothelial cell (EC) (dys)function and vascular pathophysiology remains elusive.

Thus, the present study was designed to investigate whether MICU1 deficiency or decrease is causal for atherosclerosis in mice and human. We observed that MICU1 depletion augmented inflammatory responses in human endothelial cells. In addition, EC-specific *Micu1* knockout mice (*Micu1*^ECKO^) accelerated, while EC-specific *Micu1* transgenic mice (*Micu1*^ECTg^) protected against vascular inflammation and atherosclerosis. Mechanistically, MICU1 restrains inflammation via SIRT3/SOD2/mROS pathway. Of clinical significance, two-sample Wald ratio Mendelian randomization studies revealed that increased expression of MICU1 was associated with decreased risk of coronary artery bypass grafting (CABG).

## Results

### MICU1 depletion augments inflammation response in HUVECs

To understand the role of MICU1 in endothelial function, we performed a transcriptomic profiling in HUVECs using RNA-seq after MICU1 depletion (Figure 1A). As shown in Figure 1B, downregulation of MICU1 significantly increased the expression of 494 genes while decreased that of 277 genes in HUVECs based on selection criteria of fold change ≥ 2 and FDR < 0.05. Differentially expressed genes were associated with multiple KEGG pathways linked to inflammatory responses, including TNF signaling pathway, Toll-like receptor signaling, NOD-like receptor signaling, RIG-I-like receptor signaling and JAK-STAT signaling (Figure 1C). Similarly, Gene Set Enrichment Analysis (GSEA) showed that MICU1 depletion upregulated TNF signaling and Toll-like receptor signaling pathways (Figure 1D and E). Additionally, GSEA also reveals enrichment of atherosclerosis-linked transcripts after MICU1 depletion (Figure 1F). Therefore, we hypothesized that MICU1 depletion might contribute to inflammatory responses in human endothelial cells. To test the hypothesis, we assessed the expression of inflammatory cytokines and chemokines indicated by KEGG pathway analysis. For instance, the inflammatory cytokines and chemokines including C-X-C motif ligand (CXCL) 1, CXCL2, CXCL3, C-X3-C motif chemokine ligand (CX3CL)1, interleukin (IL) 12A, IL15, C-C motif chemokine ligand (CCL) 5, vascular endothelial growth factor C (VEGFC), MYD88, colony stimulating factor (CSF) 1, interferon (IFN) β1, IL6 and CXCL10 were increased in HUVECs after MICU1 depletion compared with negative control (Figure 1G). These data were validated by qRT-PCR which confirmed that MICU1 depletion led to an activation of multiple chemokines and cytokines (Figure 1H). These results indicate that MICU1 might serve as a protective factor in controlling endothelial inflammation.

**Figure 1.**
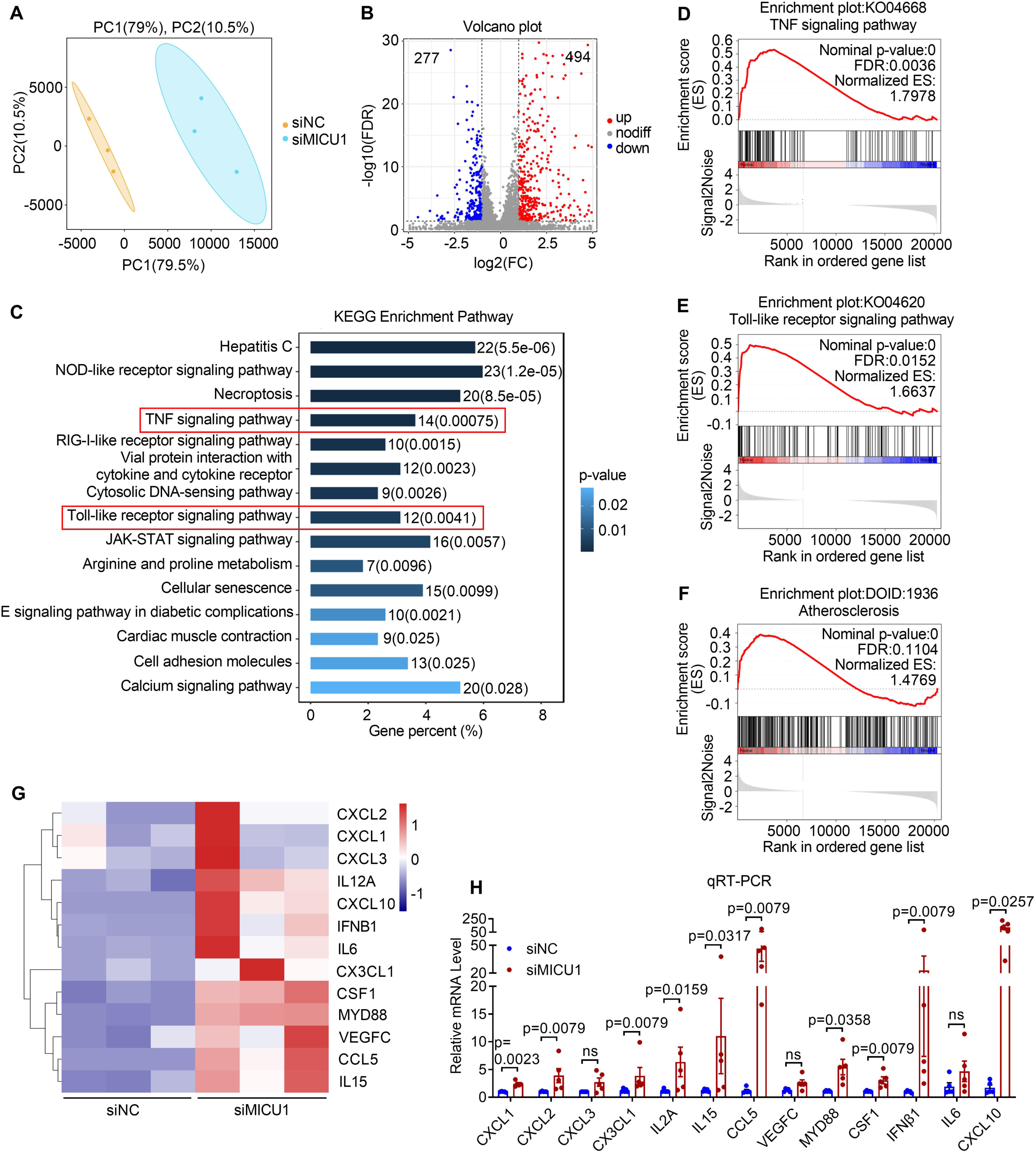
MICU1 depletion leads to an atheroprotective transcriptome profile in HUVECs. (A) Principal component analysis (PCA) comparing the transcriptomic data and plotting by co-ordinates for principal component (PC) 1 and PC2. Color-coding was used to separate treatment with negative control siRNA (siNC) or MICU1 siRNA (siMICU1). (B) Volcano plot showed differentially expressed genes after knockdown of MICU1 in human umbilical vein endothelial cells (HUVECs). Selection criteria: gene expression fold change (FC) ≥2 and false discovery rate (FDR) <0.05. (C) Kyoto Encyclopedia of Genes and Genomes (KEGG) enrichment for differentially expressed genes after MICU1 depletion. (D-F) Gene Set Enrichment Analysis (GSEA) analysis was used to examine the enrichment of TNF signaling pathway (D), Toll-like receptor signaling pathway (E) and atherosclerosis (F). (G) Heatmap showed the key differentially expressed genes from KEGG top pathway related to inflammatory chemokines or cytokines. (H) Quantitative reverse transcription polymerase chain reaction (qRT-PCR) analysis of mRNA level of inflammatory chemokines and cytokines after MICU1 depletion (n=5). Statistical analysis was performed by Welch’s *t* test (H, CXCL1, CXCL3, VEGFC, MYD88, CXCL10) and Mann-Whitney *U* test (H, CXCL2, CX3CL1, IL2A, IL15, CCL5, CSF1, IFNβ1, IL6).

### MICU1 limits inflammatory activation of endothelial cells

Based on the anti-inflammatory role of MICU1 in endothelial cells revealed by RNA-sequencing, we next explored the expression of MICU1 in response to various pro-inflammatory stimuli. As shown in Figure 2A, MICU1 protein expression was decreased in HAECs after tumor necrosis factor (TNF) α treatment, whereas the expression of MCU was slightly increased. In addition, the expression of mRNAs for MICU1 and MCU had similar changes compared with protein levels (Figure S1A and B). Besides, we detected the mRNA level of other components of the MCU complex in HUVECs, including MICU2, MICU3, MCUb and EMRE (Figure S1C-F). Furthermore, we tested MICU1 expression in HUVECs under cytokine stimulation and revealed that MICU1 was significantly decreased after the treatment with TNFα and IL-1β (Figure 2B and C). In vivo, lipopolysaccharide (LPS) was injected into mice to induce inflammation. *En face* immunofluorescence staining showed that MICU1 expression in aortic endothelium was reduced in mice treated with LPS for 6 h compared with saline treatment (Figure 2D). These results indicate the expression of MICU1 was decreased under inflammatory stimulation in vitro and in vivo.

**Figure 2.**
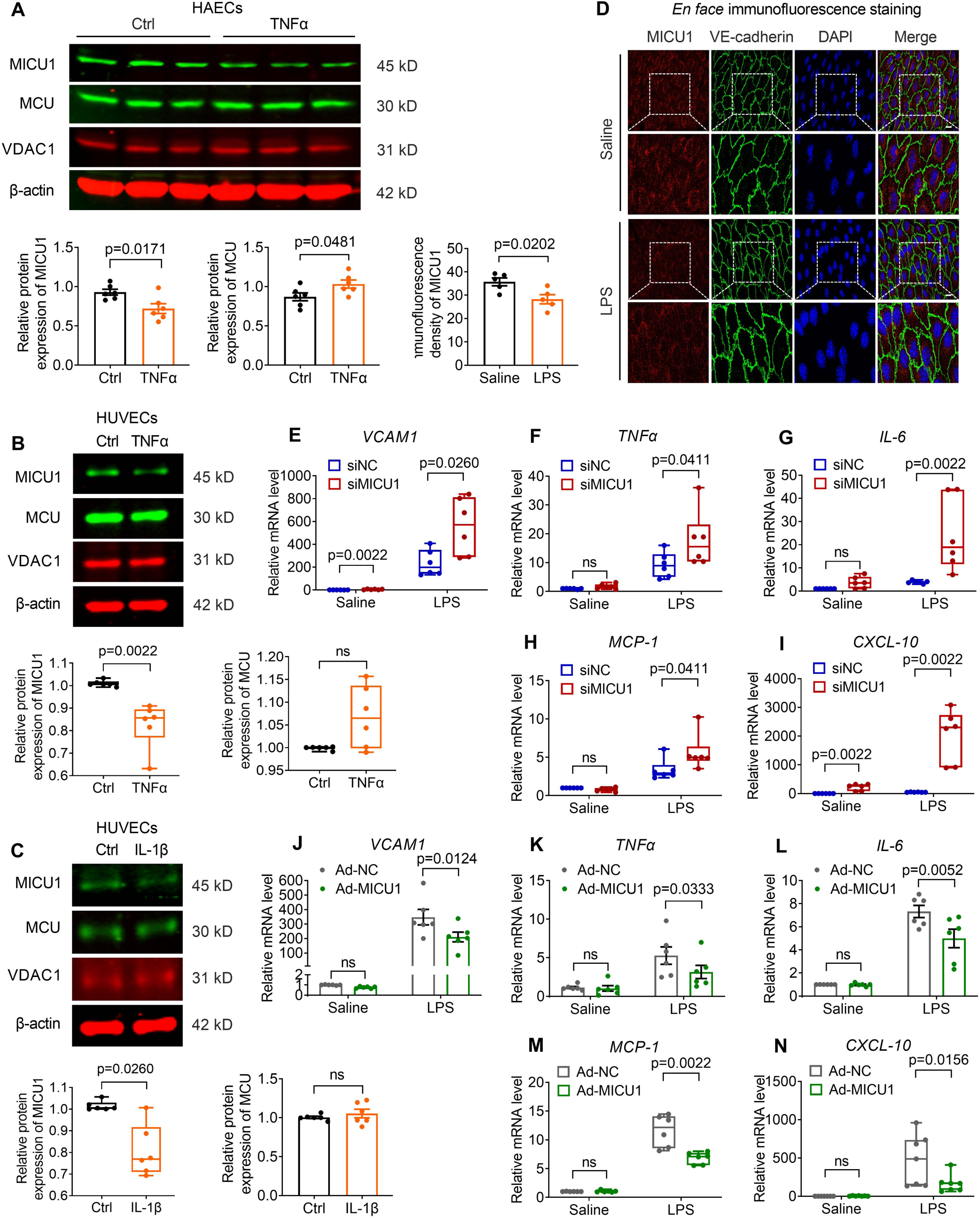
MICU1 depletion promotes endothelial inflammation. (A) Representative immunoblots and quantification data are shown for the expression of MICU1 and MCU in human aortic endothelial cells (HAECs) after treatment with tumor necrosis factor α (TNFα) (10 ng/ml) for 6 h. VADC1 and β-actin were used as the loading control (n=6). (B-C) The expression of MICU1 and MCU in human umbilical vein endothelial cells (HUVECs) exposed to tumor necrosis factor α (TNFα) (10 ng/ml) (B, n=6) or interleukin-1β (IL-1β) (10 ng/ml) (C, n=6) for 6 h. (D) *En face* immunofluorescence staining of MICU1 in mouse aortic endothelium. Mice were intraperitoneally injected with lipopolysaccharides (LPS) (10 mg/kg) for 6 h. Confocal microscopy images showed MICU1 (red), VE-cadherin (green), and DAPI (blue). Scale bars, 10 μm (n=5). (E-I) qRT-PCR analysis of mRNA level of VCAM1 (E, n=6), TNFα (F, n=6), IL-6 (G, n=6), MCP-1 (H, n=6) and CXCL-10 (I, n=6) in HUVECs after treatment with negative control siRNA (siNC) or MICU1 siRNA (siMICU1) in the presence of LPS (1 μg/ml) for 6 h. (J-N) HUVECs were transfected with negative control adenovirus (Ad-NC) or MICU1 adenovirus (Ad-MICU1) before qRT-PCR analysis of mRNA level of VCAM1 (J, n=6), TNFα (K, n=6), IL-6 (L, n=6), MCP-1 (M, n=6), and CXCL-10 (N, n=7). Ns indicates not significant. Statistical analysis was performed by Student *t* test (A, D), Mann-Whitney *U* test (B, C, left panel), Welch’s *t* test (C, right panel), multiple Mann-Whitney *U* tests (E-I, M, N) and 2-way ANOVA followed by Bonferroni post hoc tests (J-L).

The potential role of MICU1 in LPS-mediated induction of the inflammatory molecules, such as vascular cell adhesion molecule (VCAM)-1, TNFα, monocyte chemoattractant protein (MCP)-1, CXCL-10 was investigated by gene silencing. MICU1 depletion significantly elevated the expression of these molecules in the presence of LPS (Figure 2E-I), indicating that endogenous MICU1 is a negative regulator of inflammatory gene expression. In contrast, overexpression of MICU1 reduced the expression of the pro-inflammatory cytokines/chemokines (Figure 2J-N). Collectively, these results indicate that MICU1 negatively regulates endothelial inflammation.

### Endothelial MICU1 limits vascular inflammation

As Toll-like receptor (TLR) signaling pathway was enriched according to the transcriptomic profiling of MICU1 depletion, LPS, the ligand for TLR signaling, was used to investigate the role of MICU1 in endothelial inflammation in vivo (27). To examine whether MICU1 regulates inflammation in vivo, we generated mice with EC-specific ablation of *Micu1* (*Micu1*^ECKO^) by crossbreeding *Micu1*^Fl/Fl^ mice with *Cdh5*-Cre mice (Figure S2A). Successful deletion of *Micu1* in *Micu1*^ECKO^ mice was validated by genotyping and RT-PCR quantitation of *Micu1* mRNA in the aortic endothelium (Figure S2B and C). *En face* immunofluorescence staining also confirmed the ablation of endothelial MICU1 in *Micu1*^ECKO^ mice (Figure S2D). *Micu1*^ECKO^ mice have no appreciable physiological defects. *Micu1*^Fl/Fl^ littermate controls were used as wild type mice. The body weight was similar between *Micu1*^ECKO^ mice and *Micu1*^Fl/Fl^ mice (Figure S2E).

Next, mice were injected with LPS for 6 h to induce vascular inflammation. The plasma levels of the cytokines, such as IL-6, TNFα, CXCL-10, MCP-1 and E-selectin (as determined by ELISA) were remarkably increased in response to LPS treatment. Of note, *Micu1*^ECKO^ mice exhibited significantly elevated levels of IL-6, TNFα, CXCL-10 and MCP-1 after LPS treatment (Figure 3A-D), whereas the expression of E-selectin remained unaffected (Figure 3E). To further examine the effects of ablating endothelial *Micu1* on vascular inflammation, we analyzed ICAM-1 and VCAM-1 levels in aortic endothelium. Confocal microscopy showed increased ICAM-1 and VCAM-1 in the *Micu1*^ECKO^ mice endothelium compared with control mice after LPS treatment (Figure 3F and G). These findings demonstrate that *Micu1* deficiency in vascular endothelium significantly aggravated the LPS-induced inflammatory response.

**Figure 3.**
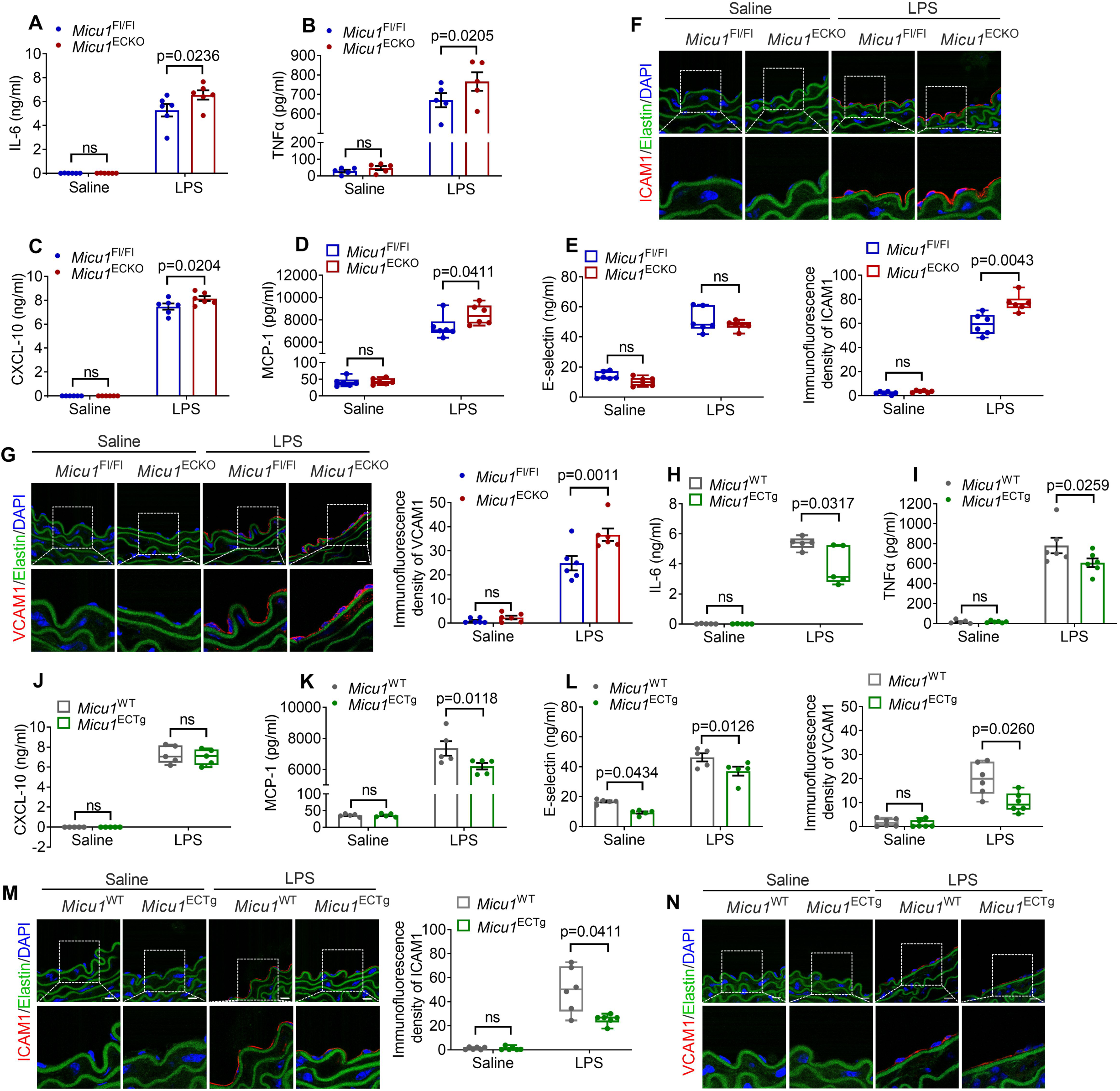
Endothelial-specific *Micu1* deficiency accelerates, while endothelial-specific *Micu1* transgenic overexpression attenuates LPS-induced vascular inflammation in mice. (A-E) Enzyme linked immunosorbent assay (ELISA) of IL-6 (A, n=6), TNFα (B, n=5), CXCL-10 (C, n=6), MCP-1 (D, n=6), E-selectin (E, n=6) in serum from *Micu1*^Fl/Fl^ mice or *Micu1*^ECKO^ mice treated with saline or LPS (10 mg/kg) for 6 h. (F-G) Representative confocal microscopy images of ICAM1 expression (F, n=6) and VCAM1 expression (G, n=6) in the aortic sections of *Micu1*^Fl/Fl^ mice or *Micu1*^ECKO^ mice exposed to saline or LPS (10 mg/kg) for 6 h. ICAM1 or VCAM1 (red), elastin (green), and DAPI (blue). Scale bars, 10 μm. (H-L) ELISA of IL-6 (H, n=5), TNFα (I, n=6), CXCL-10 (J, n=5), MCP-1 (K, n=5), E-selectin (L, n=5) in serum from *Micu1*^WT^ mice or *Micu1*^ECTg^ mice exposed to saline or LPS (10 mg/kg) for 6 h. (M-N) Representative images of ICAM1 (I, n=6) and VCAM1 protein expression (J, n=6) in the aortic sections of *Micu1*^WT^ mice or *Micu1*^ECTg^ mice exposed to saline or LPS (10 mg/kg) for 6 h. Scale bars, 10 μm. Statistical analysis was performed by 2-way ANOVA followed by Bonferroni post hoc tests (A-C, G, I, K, L) and multiple Mann-Whitney *U* tests (D-F, H, J, M, N).

Furthermore, to determine whether overexpression of MICU1 can alleviate vascular inflammation in response to LPS challenge, we used EC-specific *Micu1* transgenic mice (*Micu1*^ECTg^), which were obtained by crossbreeding *Rosa*^LSL-Micu1^ mice with *Cdh5*-Cre mice (Figure S3A). These mice were validated by tail genotyping and qRT-PCR for the mRNA level of *Micu1* in the aortic endothelium (Figure S3B and C). *En face* immunofluorescence staining showed successful overexpression of MICU1 in aortic endothelium (Figure S3D). *Micu1*^ECTg^ mice behave normally and were viable. Body weight in *Micu1*^ECTg^ mice and control mice were comparable (Figure S3E). Of note, ELISA of plasma samples revealed that LPS induction of IL-6, TNFα, MCP-1 and E-selectin was significantly attenuated in *Micu1*^ECTg^ mice compared to control mice, whereas levels of CXCL-10 were unaltered (Figure 3H-L). To validate whether overexpression of MICU1 in mice can protect vascular endothelium, we detected the expression of ICAM-1 and VCAM-1 in aortic endothelium. Immunofluorescent images showed decreased ICAM-1 and VCAM-1 expression in the endothelium of *Micu1*^ECTg^ mice compared with control mice exposed to LPS (Figure 3M and N). Taken together, these results demonstrate that *Micu1* overexpression protects vascular endothelium from LPS-induced inflammatory response in a mouse model.

### MICU1 in endothelial cells limits Western diet-induced atherosclerosis

The concerted actions of lipid deregulation and inflammatory responses lead to endothelial activation presenting with increased adhesion molecules, adhesion of leukocytes, and disrupted endothelial cell junctional stability, which may consequently accelerated atherosclerotic plaque formation (28). To explore whether MICU1 is involved in atherosclerosis in hypercholesterolemic mice, *Micu1*^ECKO^ mice and *Micu1*^Fl/Fl^ mice were treated with AAV8-PCSK9^D377Y^ concurrent with Western diet feeding for twelve weeks to induce hypercholesterolemia.

Lesion formation quantified by *en face* Oil Red O staining of aortas was significantly increased in both male and female cohorts of *Micu1*^ECKO^ mice compared with respective controls (Figure 4A). Furthermore, for male mice, Oil Red O staining (Figure 4B) and histological analysis (Figure 4C) of the aortic sinus showed that deletion of endothelial *Micu1* increased atherosclerotic plaque area. Next, several features of plaque composition were quantified. Masson staining of aortic root sections indicated that *Micu1* deficiency in endothelium reduced the proportion of collagen, a feature of plaque stability, in the aortic plaques (Figure 4D). The number of macrophages within the aortic sinus plaques was increased in *Micu1*^ECKO^ mice compared with *Micu1*^Fl/Fl^ mice, whereas α-SMA expression showed no overt differences (Figure 4E). Similar to the LPS-inflammation model, *Micu1*^ECKO^ mice also exhibited significantly elevated levels of serum IL-6, TNF-α, MCP-1 and E-Selectin compared with *Micu1*^Fl/Fl^ mice (Figure 4J-M). Thus, endothelial *Micu1* is a negative regulator of plaque growth and reduces features of plaque instability. It was concluded that the mechanism does not involve alteration in circulating levels of lipoproteins since no significant differences were observed between *Micu1*^ECKO^ mice and *Micu1*^Fl/Fl^ mice, concerning serum TG, serum CHO, serum HDL and serum LDL levels (Figure 4F-I).

**Figure 4.**
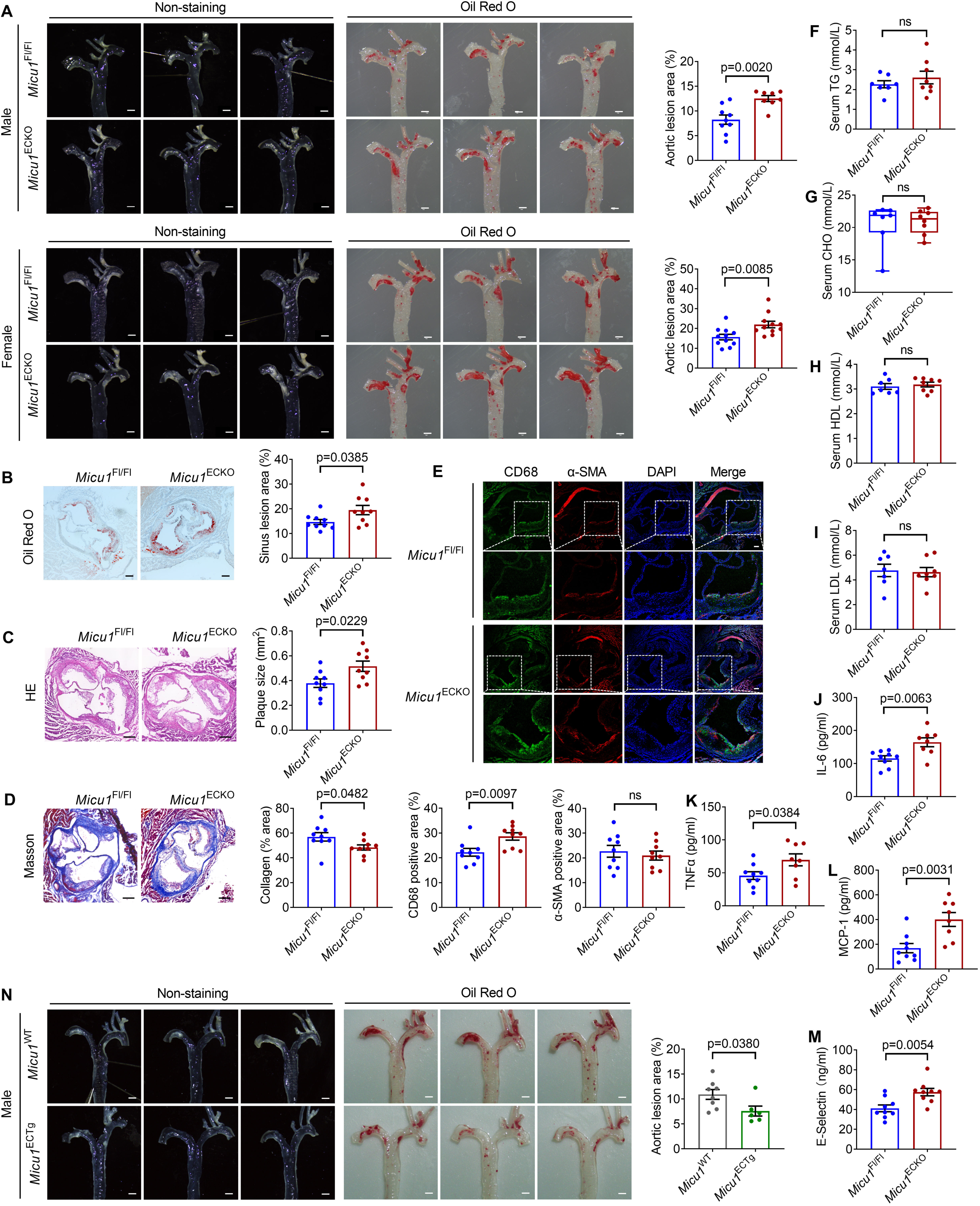
MICU1 in endothelial cells limits Western diet-induced atherosclerosis. (A) Representative images of Oil Red O staining of atherosclerotic lesions of aorta in male (n=8-9) and female (n=11) *Micu1*^Fl/Fl^ mice or *Micu1*^ECKO^ mice infected with AAV8-PCSK9^D377Y^ after 12 weeks of western diet. Scale bars, 1 mm. (B-D) Oil Red O staining (B), hematoxylin-eosin staining (H&E) staining (C) or masson staining (D) of lesions of the aortic root in male *Micu1*^Fl/Fl^ mice or *Micu1*^ECKO^ mice from (A) (n=9). Scale bars, 200 μm. (E) Staining of CD68-positive macrophages in lesion area of the aortic sinus from male *Micu1*^Fl/Fl^ mice or *Micu1*^ECKO^ mice from (A) (n=9). Scale bars, 100 μm. (F-I) Serum triglyceride (TG), cholesterol (CHO), high-density lipoprotein (HDL) and low-density lipoprotein (LDL) were detected in male *Micu1*^Fl/Fl^ mice or *Micu1*^ECKO^ mice infected with AAV8-PCSK9^D377Y^ after 12 weeks of western diet (n=7-8). (J-M) Enzyme linked immunosorbent assay (ELISA) of serum IL-6 (J), serum TNFα (K), serum MCP-1 (L), serum E-selectin (M) from male *Micu1*^Fl/Fl^ mice or *Micu1*^ECKO^ mice infected with AAV8-PCSK9^D377Y^ after 12 weeks of western diet (n=8-9). (N) Representative images of Oil Red O staining of atherosclerotic lesions of aorta in male (n=6-8) *Micu1*^WT^ mice or *Micu1*^ECTg^ mice infected with AAV8-PCSK9^D377Y^ after 12 weeks of western diet. Scale bars, 1 mm. Statistical analysis was performed by Student *t* test (A-F, H-N) and Mann-Whitney *U* test (G).

Consistent with the phenotype obtained in male mice, *Micu1*^ECKO^ female mice showed an increase of the aortic root and aortic sinus plaques, as well as infiltration of macrophages in the aortic sinus plaques compared with *Micu1*^Fl/Fl^ mice (Figure S4A, B and D). *Micu1*^ECKO^ female mice represented decreased collagen in the aortic sinus plaques (Figure S4C). No significant differences were observed between *Micu1*^ECKO^ mice and control mice, in serum lipid profile (Figure S4E-H). There was an increase of serum IL-6 and TNF-α in *Micu1*^ECKO^ mice after twelve weeks of Western diet feeding (Figure S4I and J). Serum levels of MCP-1 and E-Selectin did not change (Figure S4K and L).

As a gain-of-function approach, *Micu1*^ECTg^ mice were also used to investigate the atherosclerotic lesion formation as compared to control mice. EC-specific *Micu1* transgenic overexpression significantly decreased the plaques formation in *en face* aorta compared with control mice (Figure 4N). Collectively, these results suggest that endothelial *Micu1* limits atherosclerotic lesions in mice.

### MICU1 regulates mitochondrial Ca^2+^ uptake and ROS production in inflammed human endothelial cells

MICU1 is an essential factor responsible for mitochondrial Ca^2+^ influx regulation because its two canonical EF hands can sense Ca^2+^ and serve as the gatekeeper for MCU-mediated mitochondrial Ca^2+^ uptake (10, 16, 21). To understand the mechanisms of MICU1 in LPS-induced endothelial inflammation, we evaluated mitochondrial Ca^2+^ influx in HUVECs. Strikingly, [Ca^2+^]_m_ was elevated in response to an agonist-induced [Ca^2+^]_i_ rise in MICU1-depleted endothelial cells under basal conditions (Figure 5A), as well as in MICU1 knockdown cells under LPS treatment (Figure 5B). We therefore reasoned that overexpression of MICU1 could inhibit mitochondrial Ca^2+^ uptake in cells exposed to LPS. To test this hypothesis, HUVECs were infected with an adenovirus containing a *MICU1* cDNA overexpression cassette. Histamine-induced mitochondrial Ca^2+^ uptake rose transiently, but then showed a tendency to decrease compared with control (Figure 5C). Importantly, overexpression of MICU1 prevented elevated [Ca^2+^]_m_ induced by histamine in HUVECs exposed to LPS (Figure 5D). These results indicate that mitochondrial Ca^2+^ accumulation in cells following LPS treatment could be reduced by MICU1.

**Figure 5.**
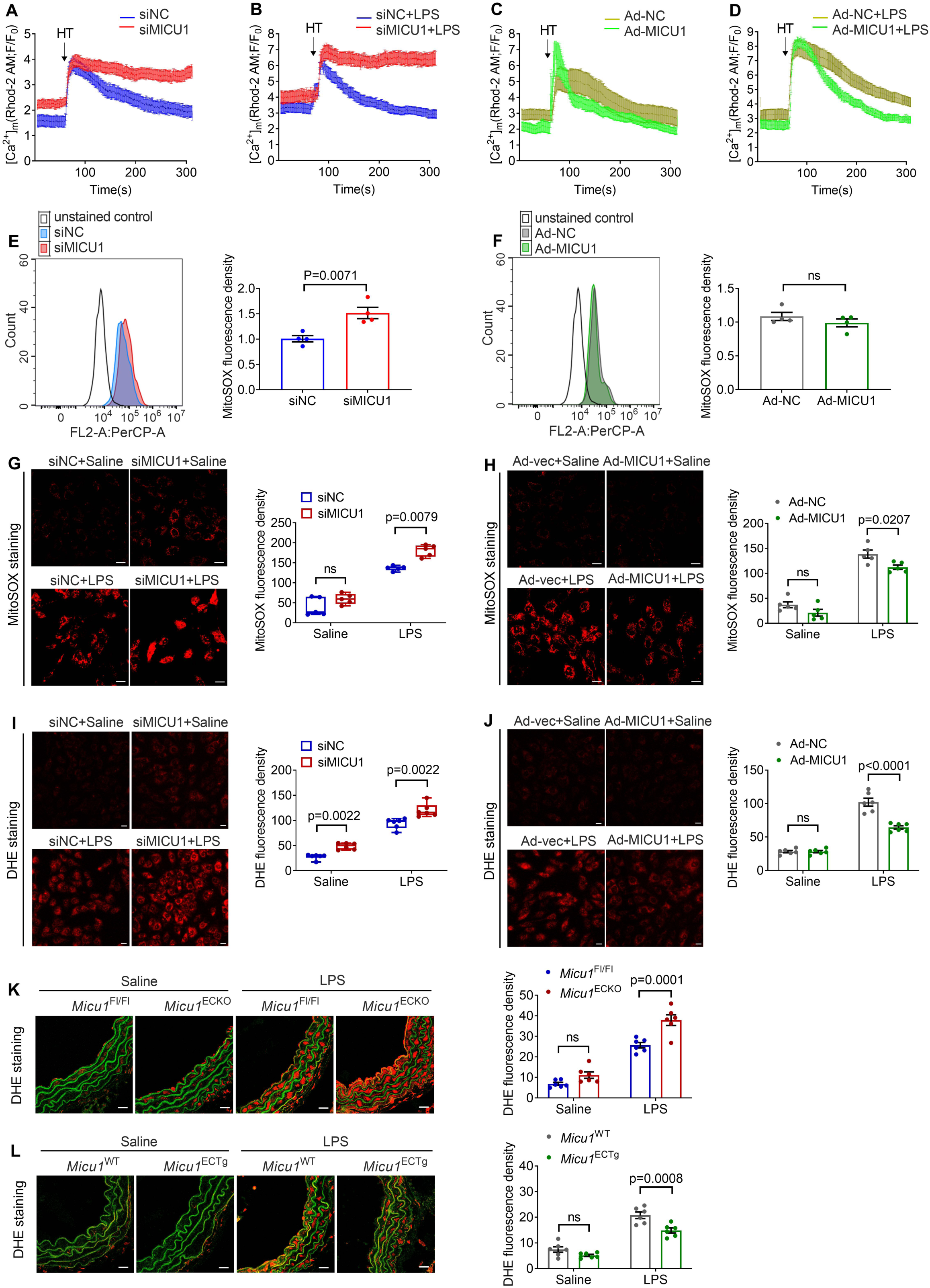
MICU1 ameliorates mitochondrial ROS via reducing mitochondrial Ca^2+^ uptake. (A-B) Kinetics of [Ca^2+^]_m_ in response to histamine (HT) (50 μM) in human umbilical vein endothelial cells (HUVECs) treated with negative control siRNA (siNC) or MICU1 siRNA (siMICU1) in the presence (B, siNC+LPS n=22, siMICU1+LPS n=16) or absence of (A, siNC n=23, siMICU1 n=16) LPS (1μg/ml) for 6 h. (C-D) Kinetics of [Ca^2+^]_m_ in response to HT (50 μM) in HUVECs treated with negative control adenovirus (Ad-NC) or MICU1 adenovirus (Ad-MICU1) in the presence (D, Ad-NC+LPS n=17, Ad-MICU1+LPS n=17) or absence of (C, Ad-NC n=13, Ad-MICU1 n=12) LPS (1μg/ml) for 6 h. (E-F) Mitochondrial ROS level was examined by mitochondrial superoxide indicator (MitoSOX) followed by flow cytometry analysis under conditions of MICU1 knockdown (E) or MICU1 overexpression (F) (n=4). (G-H) Representative images showing mitoSOX fluorescence in HUVECs under the conditions of MICU1 knockdown (G) or MICU1 overexpression (H) (n=5). Scale bars, 20 μm. (I-J) Representative images showing dihydroethidium (DHE) fluorescence in HUVECs under the conditions of MICU1 knockdown (I) or MICU1 overexpression (J) (n=6). (K) Representative images showing DHE fluorescence in aortic sections from *Micu1*^Fl/Fl^ mice or *Micu1*^ECKO^ mice treated with saline or LPS (10 mg/kg) for 6 h (n=6). Scale bars, 20 μm. (L) Representative images showing DHE fluorescence in aortic sections from *Micu1*^WT^ mice or *Micu1*^ECTg^ mice treated with saline or LPS (10 mg/kg) for 6 h (n=6). Scale bars, 20 μm. Statistical analysis was performed by Student *t* test (E, F), multiple Mann-Whitney *U* tests (G, I) and 2-way ANOVA followed by Bonferroni post hoc tests (H, J-L).

Mitochondrial Ca^2+^ uptake is required not only to match energy supply but also to maintain antioxidative capacity preventing excessive production of reactive oxygen species (ROS) (7, 22). Although a low level of ROS may have physiological roles, higher levels of ROS result in activation of kinases and inflammatory cytokines involved in cardiovascular diseases (7, 29, 30). We therefore performed flow cytometry to confirm whether the changes in [Ca^2+^]_m_ regulated by MICU1 contribute to the mitochondrial ROS (mROS) accumulation. As shown in Figure 5E, quantitative mitochondrial superoxide indicator (mitoSOX) red staining analyzed by flow cytometry indicated that MICU1 depletion led to an increased ROS concentration while overexpression of MICU1 caused small but insignificant decrease of the mitochondrial ROS (Figure 5E and F). Further, we investigated mROS level in comparison to the whole cell ROS accumulation in HUVECs in response to LPS treatment. The production of mROS was enhanced by MICU1 depletion and was abrogated by MICU1 overexpression (Figure 5G and H). The effects of MICU1 on the whole cell ROS concentration were consistent with mROS (Figure 5I and J). In vivo, EC-specific *Micu1* deletion led to excessive ROS levels of aortic sections in response to LPS-induced inflammation in mice and EC-specific *Micu1* transgenic mice showed decreased ROS levels (Figure 5K and L). These results suggest that MICU1-mediated suppression [Ca^2+^]_m_ contributes to ROS reduction in LPS-induced vascular inflammation.

### SIRT3/SOD2 pathway links MICU1 regulation of [Ca^2+^]_m_ to endothelial inflammation and Western diet-induced atherosclerosis

To define the mechanisms linking MICU1-regulated [Ca^2+^]_m_ to ROS production, we investigated the potential role of sirtuin (SIRT) 3 because this deacetylating enzyme can couple [Ca^2+^]_m_ with mitochondrial ROS by targeting SOD2 (31, 32). Since SOD2 acetylation is associated with inflammation, we next analyzed the influence of the MICU1 on the inflammatory marker VCAM-1. MICU1 silencing with or without TNFα treatment led to elevated VCAM-1 expression whereas MICU1 overexpression had the opposite effect (Figure 6A and B). It is noteworthy that SIRT3 expression was decreased in response to silencing of MICU1 in HAECs (Figure 6C). The decrease of SIRT3 in response to MICU1 silencing was exaggerated by TNFα treatment (Figure 6D) and could be reversed by MICU1 overexpression (Figure 6E and 6F). Further, we addressed the potential role of MICU1 in regulating SOD2 activity. MICU1 depletion led to an increase in acetylated SOD2 levels but did not alter total SOD2 levels (indicating SOD2 hyperacetylation) (Figure 6C and 6D), whereas overexpression of MICU1 reduced SOD2 acetylation (Figure 6E and 6F). Moreover, the effect of MICU1 on SOD2 activity was confirmed by SOD2 activity detection kit (Figure 6G and H).

**Figure 6.**
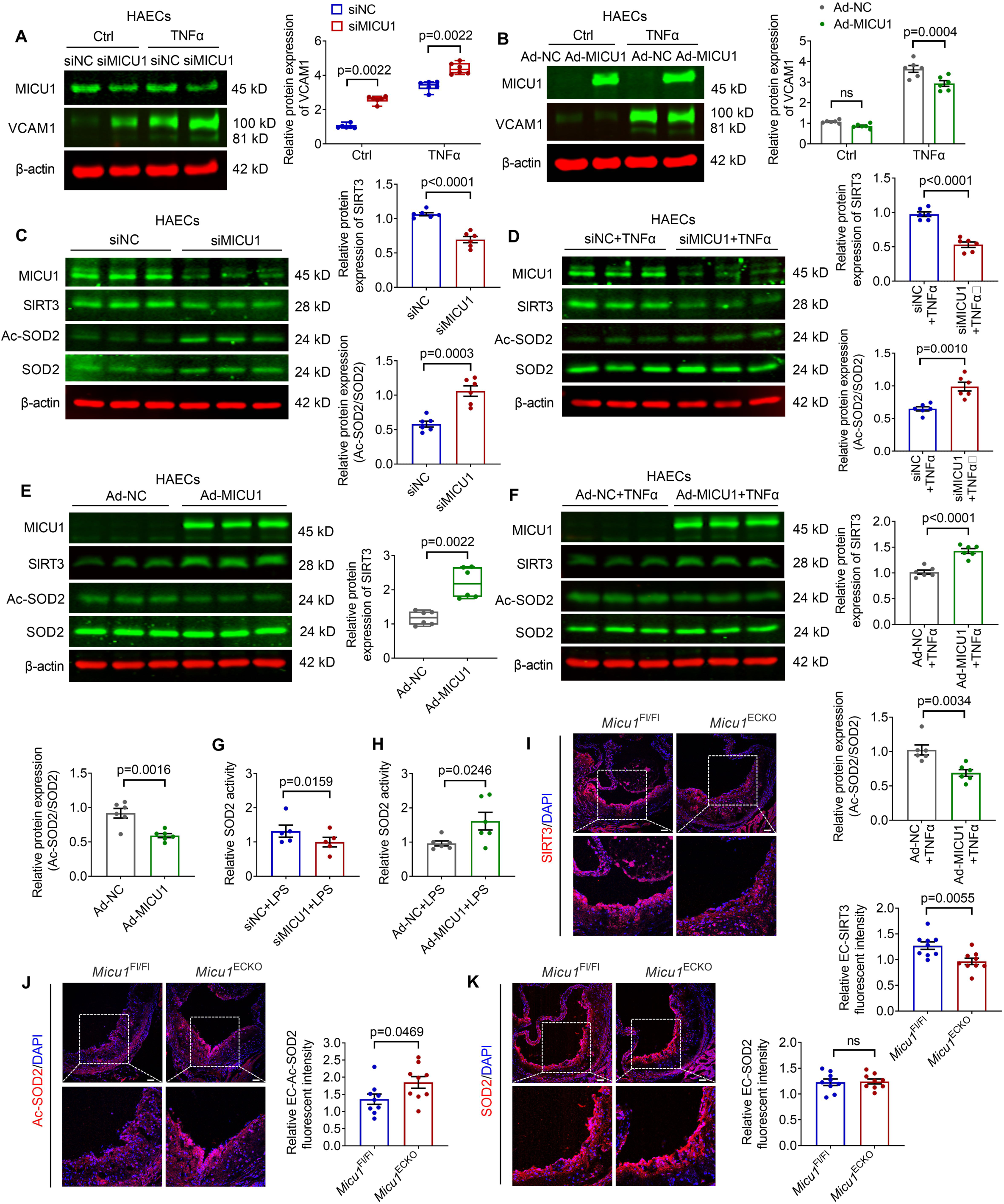
SIRT3/SOD2 pathway is involved in MICU1-mediated regulatory effects on [Ca^2+^]_m_, endothelial inflammation and diet-induced atherosclerosis. (A) The expression of VCAM1 was measured by immunoblot in human aortic endothelial cells (HAECs). Cells were treated with negative control siRNA (siNC) or MICU1 siRNA (siMICU1), and then exposed to TNFα (10 ng/ml) for 6 h (n=6). (B) The expression of VCAM1 was measured by immunoblot in HAECs. Cells were treated with negative control adenovirus (Ad-NC) or MICU1 adenovirus (Ad-MICU1) and then exposed to TNFα (10 ng/ml) for 6 h (n=6). (C-F) Protein expression of SIRT3 and Ac-SOD2 (acetylated SOD2) were determined by immunoblot after MICU1 depletion (C) or overexpression MICU1(E) (n=6). Expression of SIRT3, SOD2 and Ac-SOD2 were determined by immunoblot after MICU1 depletion (D) or overexpression MICU1(F) in HAECs treated with TNFα (10 ng/ml) for 6 h (n=6). (G-H) SOD2 enzyme activity was assayed in HUVECs after MICU1 depletion (G) or MICU1 overexpression (H) (n=5). (I-K) Immunofluorescence staining of SIRT3 (I), Ac-SOD2 (J), SOD2 (K) in lesion area of the aortic sinus sections from male *Micu1*^Fl/Fl^ mice or *Micu1*^ECKO^ mice infected with AAV8-PCSK9^D377Y^ after 12 weeks of western diet (n=9). Scale bars, 50 μm. Statistical analysis was performed by multiple Mann-Whitney *U* tests (A), 2-way ANOVA followed by Bonferroni post hoc tests (B), Student *t* test (C, D, E, Ac-SOD2/SOD2, F-K) and Mann-Whitney *U* test (E, SIRT3).

To validate the potential role of SIRT3/SOD2 pathway in vivo, we analyzed SIRT3 expression and levels of acetylated SOD2 in aortic root sections of hypercholesterolemic *Micu1*^ECKO^ and *Micu1*^Fl/Fl^ mice. *Micu1*^ECKO^ mice showed a decrease in SIRT3 expression and an increase in Ac-SOD2 level compared with *Micu1*^Fl/Fl^ mice (Figure 6I-K), suggesting that *Micu1* enhances SIRT3 and reduces SOD2 acetylation. Taken together, EC-specific *Micu1* deletion links Western diet-induced atherosclerosis to SIRT3/SOD2 pathway.

### SIRT3 depletion blocks the function of MICU1

To further examine the role of SIRT3 in vascular endothelial dysfunction caused by MICU1 depletion, we silenced the expression of SIRT3 in HUVECs (Figure S5). As described above, upregulation of MICU1 resulted in decreased mitochondrial ROS level, while SIRT3 knockdown blunted the effect of MICU1 overexpression in both HUVECs and HAECs (Figure 7A and B). The elevated mROS caused by MICU1 depletion was scavenged by mitoTEMPO (Figure 7C). Western blot analysis showed reduced SOD2 activity and vascular inflammation markers of VCAM-1 inhibited by MICU1 overexpression also were reversed by SIRT3 depletion (Figure 7D). These data suggest a critical role of SIRT3 in mediating the protective roles of MICU1 in vitro.

**Figure 7.**
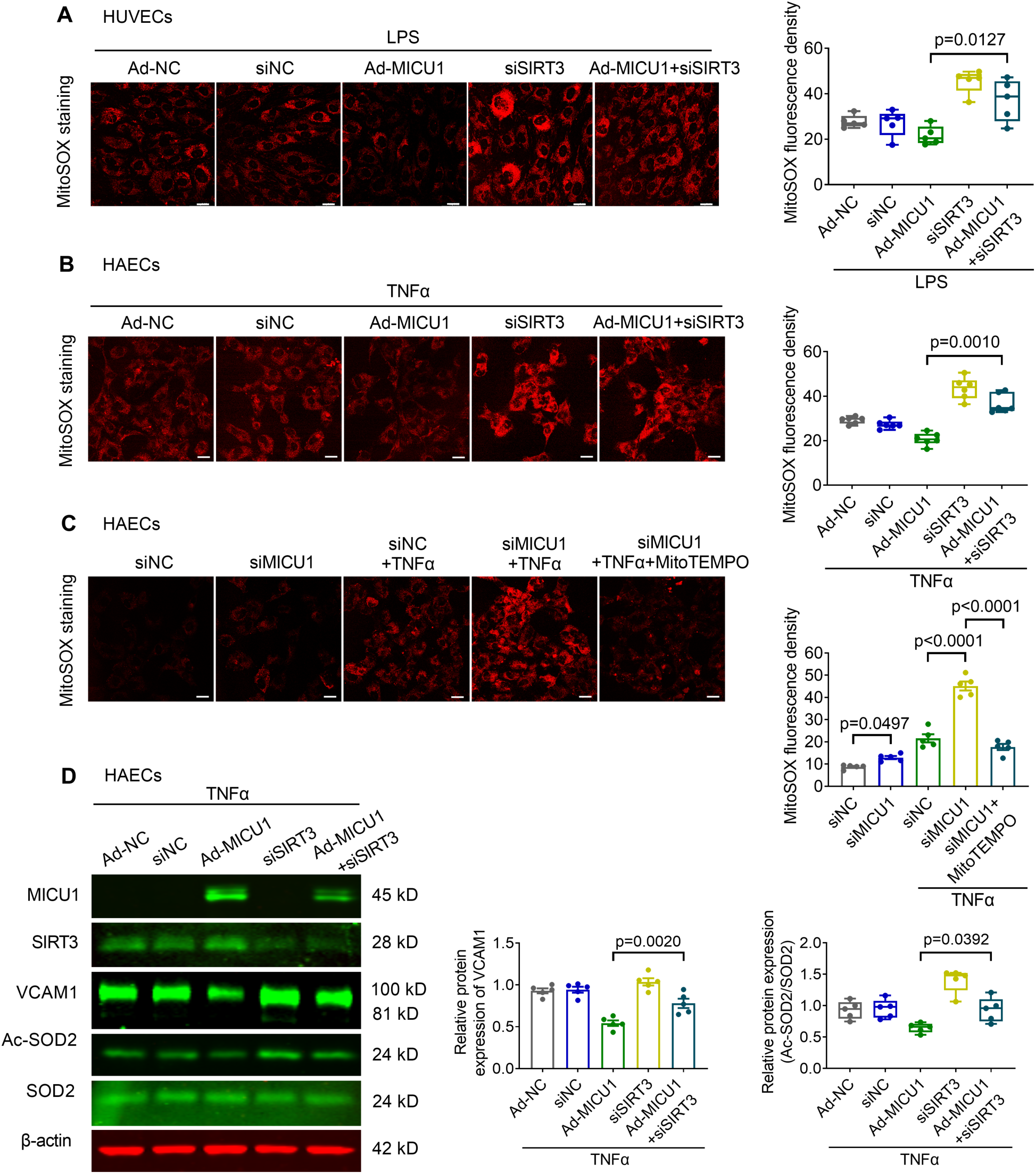
SIRT3 depletion blocks the effects of MICU1 on endothelial dysfunction. (A) Representative images showing mitochondrial superoxide indicator (MitoSOX) fluorescence in human umbilical vein endothelial cells (HUVECs) with different treatment as described (n=5). (B) Representative images showing mitochondrial superoxide indicator (MitoSOX) fluorescence in human aortic endothelial cells (HAECs) with different treatment as described (n=6). (C) Representative images showing mitochondrial superoxide indicator (MitoSOX) fluorescence in HAECs transfected with negative control siRNA (siNC) or MICU1 siRNA (siMICU1), and then exposed to TNFα (10 ng/ml). MitoTEMPO (5 μM) was added for 1 h (n=5). (D) Protein expression of Ac-SOD2 and VCAM1 in HAECs were determined by immunoblot with different treatment as described (n=5). Statistical analysis was performed by Kruskal-Wallis test followed by Dunn’s multiple comparisons test (A, B, D, right panel) and 1-way ANOVA followed by Bonferroni post hoc tests (C, D, left panel).

### Association of vascular MICU1 expression with CABG in humans

To explore the role of endothelial MICU1 gene expression in human cardiovascular diseases, we utilized Mendelian randomization (MR) studies. We constructed genetic instrumental variables (IVs) that represent changes in MICU1 gene levels in blood vessels, using genetic variant associated with MICU1 gene expression in tibial artery tissue sourced from GTEx. Two-sample Wald ratio MR revealed that increased expression of MICU1 was associated with decreased risk of CABG (FinnGen) (Odds ratio [OR] = 0.78, 95% CI, 0.67 to 0.92, P =0.003) (Figure 8A). The sensitivity analysis, which focused on coronary artery tissue, yielded similar results for CABG (OR = 0.81, 95% CI, 0.71 to 0.94, P =0.004). An independent MR analysis using CABG data in UKB, which involved significantly fewer cases, yielded a very similar effect size (OR = 0.77) with a suggestively significant result (P = 0.07). We then combined the two CABG summary datasets to enhance the statistical power and obtain more precise estimations (OR = 0.78, 95% CI, 0.68 to 0.90, P = 0.0006). Furthermore, analysis of eQTL data revealed that the risk allele (C) associated with CABG inversely correlates with MICU1 gene expression in both tibial and coronary artery tissues (Figure 8B). This is consistent with the study’s findings that MICU1 gene expression levels are decreased in vascular plaque tissues. In contrast, we did not detect associations between expression of MICU1 gene and other cardiovascular events (Figure 8A). In an additional sensitivity analysis, we conducted Bayesian enumeration colocalization to assess the presence of a shared causal variant that influences both MICU1 gene expression and CABG. We found intermediate-to-strong evidence supporting this shared causal variant (Figure 8C and 8D). We conclude that vascular expression of MICU1 in humans correlates inversely with risk of CABG in humans.

**Figure 8.**
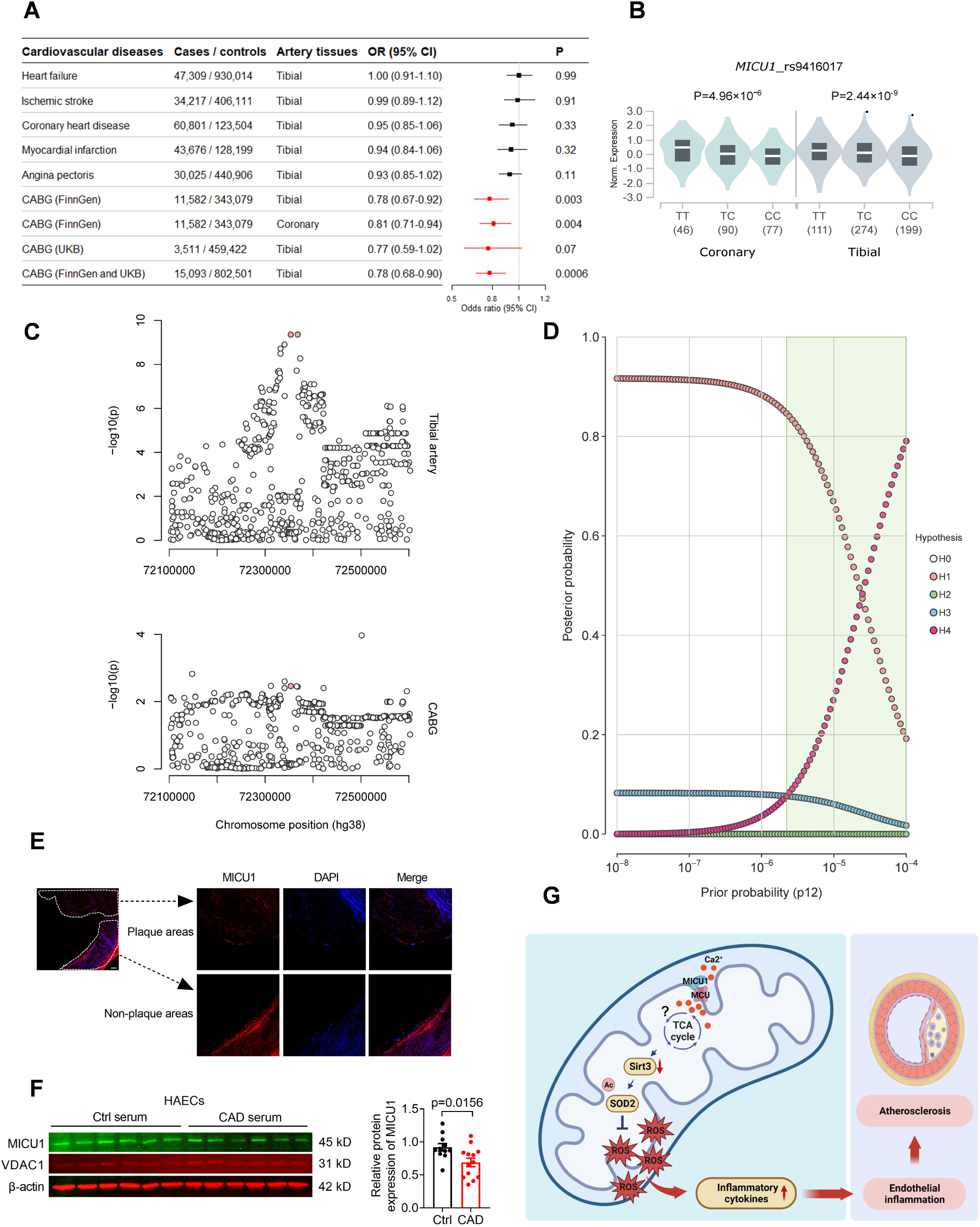
Association of vascular MICU1 expression with CABG in humans. (A) Results of 2-sample Wald ratio Mendelian randomization, testing the effects of expression of *MICU1* gene in vascular artery tissues on cardiovascular diseases. (B) Analysis of eQTL data from the GTEx database (https://www.gtexportal.org/home/) revealed that the risk allele (C) associated with coronary artery bypass grafting (CABG) exhibits an inverse relationship with *MICU1* gene expression in both tibial and coronary artery tissues for a lead eQTL (rs9416017). (C) Regional association plots highlighting ± 250 kb surrounding the lead eQTL in *MICU1* locus for tibial artery (top) and coronary artery bypass grafting (CABG) from the FinnGen study on chromosome 10. (D) Results from a Bayesian enumeration colocalization sensitivity analysis are presented, where each hypothesis represents a distinct causal setup. For a spectrum of prior probabilities (default P_12_ = 1 × 10^-5^), the posterior probability for each hypothesis is depicted in a plot. H0, neither trait has a genetic association in the region; H1, only *MICU1* expression in tibial artery has a genetic association in the region; H2, only CABG has a genetic association in the region; H3, both traits are genetically linked, but distinct causal variants; and H4, both traits have genetic associations and share a single causal variant. From the default (P_12_ = 1 × 10^-5^) to more optimistic (P_12_ = 1 × 10^-4^) priors (corresponding to approximately 0.8% and 8.0% probabilities of a shared causal variant), there is intermediate (27.4%) to strong (79.0%) posterior probability for a shared causal variant at the *MICU1* locus surrounding the lead eQTL. The shaded green area represents the range of prior probabilities for which H4 is more probable than H3. (E) Representative immunofluorescence is shown for the expression of MICU1 in human aortas with atherosclerosis. Confocal microscopy images showed MICU1 (red), and DAPI (blue). Scale bars, 100 μm (n=4). (F) The expression of MICU1 in human aortic endothelial cells (HAECs) treatment with human serum of coronary artery disease compared with healthy condition for 24 h (n=11-13). Statistical analysis was performed by Student *t* test. (G) Working model: MICU1 serves as a gatekeeper of mitochondria Ca^2+^ uniporter, which maintains mitochondrial Ca^2+^ balance. MICU1 deficiency causes mitochondria Ca^2+^ overload followed decreased SIRT3, increased Ac-SOD2 and inhibited SOD2 activity. Therefore, mitochondrial reactive oxygen species (mROS) are excessively accumulated due to impaired SOD2-mediated mROS scavenging, resulting in inflammation cytokine expression, endothelial inflammation and atherosclerosis.

To further demonstrate whether MICU1 expression was decreased in vascular endothelium from atherosclerotic tissues, we collected the atherosclerotic aortas from patients with coronary artery disease who underwent aortic surgery, and detected MICU1 level in plaque region. The expression of MICU1 was decreased in the intima of human aortas with atherosclerotic plaques compared with non-plaque region (Figure 8E). Besides, the MICU1 level was decreased in HAECs after treatment with serum from patients of coronary artery disease (CAD) compared with healthy subjects (Figure 8F). These data indicate a possible relevance between MICU1 level and atherosclerosis.

## Discussion

In the present study, we revealed that MICU1 is a crucial factor in regulating endothelial inflammation and consequently preventing the development of atherosclerosis, as summarized in Figure 8G. MICU1 depletion triggered inflammation in endothelial cells. Moreover, *Micu1* deletion aggravated vascular inflammation and atherosclerosis, whereas transgenic overexpression of *Micu1* prevented vascular inflammatory responses and atherosclerosis in mice. Mechanistically, the downregulation of MICU1 contributed to an excessive accumulation of mitochondrial Ca^2+^, promoting ROS production via decreasing the SIRT3 expression followed by hyperacetylation of SOD2. This study uncovers a novel role of endothelial MICU1 in vascular inflammation and atherosclerosis.

MICU1 regulates bioenergetics, mitochondrial cristae structure and cell function, thus serving as a gatekeeper of MCU to regulate mitochondrial Ca^2+^ uptake (21, 23, 33, 34). Recent studies have implicated MICU1 in regulating not only mitochondrial Ca^2+^ uptake but also Na^+^ influx when its EF-hands are not occupied (35, 36). For the complex and uncertain mechanism described in vitro, we demonstrate the function of MICU1 also in vivo to confirm whether MICU1 deficiency leads to vascular inflammation and atherosclerosis. Using genetically modified animals, our data strongly support the concept that endothelial *Micu1* deficiency aggravates vascular inflammation and atherosclerosis.

MICU1 were found to participate in neurodegenerative diseases and muscle disorders (26, 37–39). Patients carrying MICU1 mutations have been reported developmental delay and early onset myopathy, attention deficits, insomnia and impaired cognitive pain perception (38). Recently, MICU1 has also been found related to diabetic cardiomyopathy (40). Interestingly, Patel et al. reported that MICU1 level was altered in endothelial cells under different shear stress, indicating a tight regulation of MICU1 on physiological and pathological hemodynamic (41). The modulating effects of MICU1 on [Ca^2+^]_m_ level influences not only mitochondrial bioenergetics but also the level of mROS which is closely related to vascular inflammation (6, 7, 21). However, the function of MICU1 in the vascular diseases remain largely elusive because of the lack of support from genetic animal models. In the present study, we generated EC-specific *Micu1* deficient mice and EC-specific *Micu1* transgenic mice to explore the effects of MICU1 in vivo. We found that the expression of MICU1 was decreased after treatment with LPS, TNFα and IL-1β in endothelial cells, as well as in LPS-treated mice. Moreover, the LPS-induced vascular inflammatory response was aggravated in *Micu1*^ECKO^ mice, while it was relieved in *Micu1*^ECTg^ mice.

Proinflammatory cytokines and chemokines amplify the development of atherosclerosis in animal models (28). Blocking inflammation with an anti-IL-1β antibody ameliorated cardiovascular disease outcomes in a trial of Canakinumab Anti-inflammatory Thrombosis Outcome Study (42, 43). Further, in a recent analysis of 13970 contemporary high-risk patients with statin intolerance revealed that inflammation assessed by high-sensitivity C-reactive protein (hsCRP) was a stronger predictor of future risk for cardiovascular events than hyperlipidemia assessed by low-density importance of inflammation for predicting cardiovascular death (4). As the effects of MICU1 on regulating inflammatory responses, we then observed the role of MICU1 on Western diet-induced atherosclerosis. *Micu1*^ECKO^ mice exhibited an increase in atherosclerotic plaque size in the aortic root in both male and female mice and the atherosclerotic plaque tended to be unstable. However, overexpression of MICU1 in endothelial cells protected against atherosclerosis. Deletion of endothelial *Micu1* increased the macrophage infiltration in the atherosclerotic plaque and elevated the plasma levels of IL-6, TNFα, MCP-1 and E-Selectin in atherosclerotic mice. The increase in features of plaque instability might be due to the increased inflammatory cytokines and chemokines, as the levels of circulating lipoproteins were unaltered by endothelial *Micu1* deletion.

Mitochondrial Ca^2+^ level was closely related to mROS, which is involved in multiple physiological and pathological conditions especially inflammatory response (8, 9). Considering previous reports demonstrating the role of MICU1 in mROS generation through its modulation of [Ca^2+^]_m_ (21, 44), we examined the role of MICU1 in the [Ca^2+^]_m_ and mROS level in MICU1 depleted endothelial cells and *Micu1*^ECKO^ mice. Consistent with these reports, our data showed downregulation of MICU1 promoted mitochondrial Ca^2+^ influx and mROS accumulation with or without LPS treatment. Furthermore, *Micu1*^ECKO^ mice with LPS treatment markedly increased ROS level in aortas. These findings suggested that MICU1 regulated vascular inflammation partially by modulating [Ca^2+^]_m_ and mROS levels.

The mROS production and ensuing mitochondrial dysfunction are partially regulated by mitochondria-localized sirtuin 3 (SIRT3). Decreased SIRT3 expression by cardiovascular risk factors is involved in disrupting mitochondrial homeostasis (31, 45–47). SIRT3 downregulation plays a causative role in vascular dysfunction and inflammatory response (31, 48, 49). It has been reported that excessive mitochondrial Ca^2+^ influx inhibited NAD^+^-dependent deacetylase activity of SIRT3, followed by a decrease in SOD2 activity, which contributes to aberrant ROS production and metastasis of hepatocellular carcinoma (32). In the present study, our data demonstrated that the downregulation of MICU1 reduced SIRT3 expression both in cultured endothelial cells and atherosclerotic lesions in a mouse model. SIRT3, as a NAD^+^-dependent mitochondrial deacetylase, positively regulates SOD2 activity by deacetylation of specific lysine residues. Furthermore, SOD2-K68 acetylation is markedly increased in Sirt3 knockout mice, which is prohibited in Sirt3 transgenic mice (31). This leads to the enhancement of endothelial permeability, activation of inflammasome pathways and vascular inflammation. Downregulation of SIRT3 in endothelial progenitor cells (EPC) also inhibits the re-endothelialization capacity of EPC, while SOD2 mimic prevents mitochondrial damage and improves EPC function (50). In addition, SOD2 deficiency in mice has already been proven to promote atherosclerotic lesion development in apolipoprotein E^-/-^ mice (51). Thus, we hypothesize that MICU1 regulates vascular inflammation and atherosclerosis potentially through SIRT3/SOD2/mROS pathway.

Our data further suggest that MICU1 regulated vascular inflammation by reducing SIRT3 level. Indeed, silencing of SIRT3 blocked the effect of MICU1 overexpression on SOD2 acetylation, mROS production and endothelial inflammation. It is well-established that SIRT3 is NAD^+^-dependent mitochondrial deacetylase closely relevant to the NAD^+^/NADH ratio. Previous studies have shown that mitochondrial Ca^2+^ triggers the activation of Ca^2+^-sensitive dehydrogenases of the tricarboxylic acid cycle, resulting in increased NADH production. Therefore, mitochondrial Ca^2+^ overload attenuates SIRT3 deacetylases activity to inhibit the activity of SOD2, a mitochondrial antioxidant enzyme, resulting in impaired mROS scavenging capacity. Asimakis *et al*. reported that SOD2 is a critical determinant in resisting oxidative stress in the heart based on genetic mouse data (52). Dikalova et al. have reported that cardiovascular risk factors reduce the expression of SIRT3, thus contributing to SOD2 hyperacetylation, which leads to mROS generation and related vascular disorders such as vascular inflammation and hypertension (31). In this study, we demonstrated the role of MICU1 in SIRT3/SOD2 activity to explore its involvement in the pathogenesis of vascular inflammation and atherosclerosis. Intriguingly, Winnik *et al*. reported that SIRT3 depletion does not affect atherosclerosis but impairs rapid metabolic adaptation and increases systemic oxidative stress in low-density lipoprotein receptor knockout mice (53). This could be due to the fact that the global knockout of SIRT3 may yield mixed results beyond regulating endothelial dysfunction. To elucidate the crucial role of endothelial cell SIRT3, EC-specific *Sirt3* knockout mice might be necessary. Indeed, Cao et al. recently reported that *Sirt3*^ECKO^ enhanced plaque formation during atherosclerosis in mice (54), which support our study that MICU1 deficiency promoting atherosclerosis may through SIRT3/SOD2 pathway. However, to elucidate the crucial role of SIRT3, the rescue experiments aimed at MICU1 and SIRT3 might be necessary in the future work.

Taken together, the present study reveals the novel role of MICU1 in regulating endothelial dysfunction and atherosclerosis. MICU1 deficiency causes mitochondrial Ca^2+^ overload followed by inhibition of SIRT3/SOD2 pathway to accumulate excessive mROS and thus activating vascular inflammation and accelerating atherosclerosis. MICU1 thus represents a novel promising therapeutic target that might be explored to manage atherosclerotic cardiovascular disease.

## Materials and methods

The data and study materials related to this study are available from the corresponding author upon reasonable request. Detailed procedures are provided in the Supplemental Material.

### Animals

*Micu1* flox mice (*Micu1*^Fl/+^) and *Cdh5*-Cre mice were generated on C57BL/6J background by Shanghai Model Organisms Center (Shanghai, China). EC-specific *Micu1* knockout mice (*Micu1*^ECKO^) were constructed by crossbreeding *Micu1*^Fl/Fl^ mice with *Cdh5*-Cre mice. EC-specific *Micu1* transgenic mice (*Micu1*^ECTg^), which were obtained by crossbreeding Rosa26-CAG-LSL-Micu1-WPRE-polyA (*Rosa26*^LSL-Micu1^) mice (Shanghai Model Organisms, Shanghai, China) with *Cdh5*-Cre mice. All animal procedures used in this study were approved by the animal ethics committee of University of Science and Technology (No. USTCACUC212301048) and in accordance with the guidelines by the Institutional Animal Care and Use Committee.

### Human tissue samples

The collection of human aortic samples or serum were approved by the institutional review board (IRB) of The First Affiliated Hospital of University of Science and Technology (No. 2021 KY 089). All experiments were performed in accordance with the relevant guidelines and regulations. Human tissues and serum were all collected from The First Affiliated Hospital of University of Science and Technology of China. Human aortic samples were obtained from patients with coronary artery disease who underwent aortic surgery. Serum samples of healthy people were from physical examination center of The First Affiliated Hospital of University of Science and Technology of China, and serum samples of people with coronary artery disease were collected from vascular surgery.

### Mendelian randomization and colocalization

Two-sample Mendelian randomization (MR) using summary statistics was performed using the TwoSampleMR (55) package (version 0.5.7) in R (version 4.3.2). Genetic instruments were constructed using conditionally independent expression quantitative trait loci (eQTL) for *MICU1* gene in vascular tissues from GTEx (version 8) (56, 57). The significant instrumental variable (IV) meeting a stringent threshold (P < 5 × 10^−8^) was found only in tibial artery tissue. We also assessed the robustness of the MR findings using a more lenient threshold (P < 5 × 10^−6^) in coronary artery tissue as part of sensitivity analysis. Corresponding effects for selected IV on ischemic stroke (58), coronary heart disease (59), myocardial infarction (59), heart failure (60) and angina pectoris(61) were obtained from the MRC IEU OpenGWAS database (https://gwas.mrcieu.ac.uk/) via the MR-Base platform (55).

The largest dataset of coronary artery bypass grafting (CABG) cases was obtained from the FinnGen study (R10) (62). We also obtained summary statistics for CABG from the UK Biobank (UKB) (63) through the MRC IEU OpenGWAS database (https://gwas.mrcieu.ac.uk/) (55). The UKB data analysis was done to validate the MR findings for CABG from the FinnGen study. We then employed an inverse-variance-weighted fixed-effects model meta-analysis to combine the two CABG summary data for enhancing the statistical power. After harmonization of effect alleles, MR was performed using the Wald ratio method (when a single variant associated with gene expression was present) or the inverse variance weighted method with multiplicative random effects (when multiple genetic IVs associated with *MICU1* expression were present). P values of less than 0.05 were considered significant.

Bayesian enumeration colocalization was also performed as a sensitivity analysis for significant associations using the coloc package in R (version 4.3.2) (64, 65). Genetic variants ± 250 kb surrounding a lead eQTL in the *MICU1* locus were obtained from GTEx (version 8), with summary statistics deposited in the eQTL catalog and FinnGen study (R10) (56, 57, 62). Given the sensitivity of enumeration colocalization to the Bayesian priors specified, we explored a spectrum of priors that reflected different anticipated probabilities of a shared causal variant affecting both *MICU1* gene expression and CABG. We considered both the default prior (P_12_ = 1 × 10^-5^) and a range of more optimistic priors (66).

### Resource availability

The data for transcriptome analysis are available in the Genome Sequence Archive in National Genomics Data Center, China National Center for Bioinformation / Beijing Institute of Genomics, Chinese Academy of Sciences (GSA-Human: HRA005700, https://ngdc.cncb.ac.cn/gsa-human).

### Statistical analysis

Statistical analyses were performed by GraphPad Prism 9.0 software or SPSS 24.0 software. All parametric data are presented as mean ± SEM. The data distribution was verified by Shapiro-Wilk normality test and the homogeneity of variance was detected by the Brown-Forsythe test. For comparison between two groups, the unpaired Student *t* test was used for data of normally distribution with equal variances, the Welch’s *t* test was used for data of normal distribution but unequal variances and the Mann-Whitney test was used for data of nonnormal distribution. For groups of three or more, 1-way analysis of variance (ANOVA) followed by Bonferroni post hoc tests was used for data of normal distribution, the Kruskal-Wallis test followed by Dunn’s multiple comparisons test was used for data of nonnormal distribution. For grouped analysis, 2-way ANOVA followed by Bonferroni post hoc tests was used for data of normal distribution and multiple Mann-Whitney tests was used for data of nonnormal distribution.

## Author contributions

Conceptualization: S.X., J.W. Methodology: L.S., R. L., M.S., Z.Z., Z.W., H.J. Investigation: L.S., R. L., Q.H., M.S., Z.Z., Z.L., M.L. Writing original draft: L.S., S.X., J.W. Review & editing: S.X., J.W., L.W., Y.H., P.C.E., J.P., G.G.C., B.C.B., S.O., J.G. All authors reviewed the manuscript.

## Acknowledgements

We want to acknowledge the participants and investigators of the FinnGen study. This study was supported by grants from the National Key R&D Program of China (Grant No. 2021YFC2500500), the National Natural Science Foundation of China (Grant Nos. 82003741, 82070464, 82370444) and the Strategic Priority Research Program of the Chinese Academy of Sciences (Grant No. XDB38010100). This work was also supported by the Program for Innovative Research Team of The First Affiliated Hospital of USTC (CXGG02), Anhui Provincial Key Research and Development Program (Grant No. 202104j07020051), Anhui Provincial Natural Science Foundation (Grant No. 2208085J08). P.C.E. was supported by the British Heart Foundation. S.X. is a recipient of Humboldt Research Fellowship from Alexander von Humboldt Foundation, Germany.

