## Supplemental Files for "Endothelial MICU1 protects against vascular inflammation and atherosclerosis by inhibiting mitochondrial calcium uptake"

Lu Sun, *et al*.

Author for correspondence:

Jianping Weng, MD,

Suowen Xu, PhD,

**The PDF file includes:**

- Supplemental methods
- Supplemental tables
- Supplemental figures

### Supplemental Methods

#### Animals

All animal procedures used in this study were approved by the animal ethics committee of University of Science and Technology of China (No. USTCACUC212301048) and in accordance with the guidelines by the Institutional Animal Care and Use Committee. Mice were housed in the temperature maintained at 22℃ with a 12-hour light/dark cycle. Mice were allowed free access to water and food unless otherwise indicated.

#### Generation of endothelial cells-specific *Micu1* knockout mice

*Micu1* flox mice (*Micu1*$^{Fl/+}$) and *Cdh5*-Cre mice (NM-KI-200173) were generated on C57BL/6J background by Shanghai Model Organisms Center (Shanghai, China). *Micu1* flox mice (*Micu1*$^{Fl/+}$) were established by CRISPR/Cas9 technology at chromosome 10 in C57BL/6J background. Guide-RNA, Cas9 mRNA, and a vector containing donor DNA were injected into fertilized eggs by microinjection. Positive mice containing *Micu1* flanked by loxP sites (Flox) were selected from offspring by sequencing. The Cre/LoxP strategy was used to generate EC-specific *Micu1* knockout mice (*Micu1*$^{ECKO}$). Briefly, *Micu1*$^{Fl/+}$ mice were further intercrossed to obtain *Micu1*$^{Fl/Fl}$ mice. *Micu1*$^{ECKO}$ were constructed by crossbreeding *Micu1*$^{Fl/Fl}$ mice with *Cdh5*-Cre mice, and *Micu1*$^{Fl/Fl}$ mice were used for controls. The identification of *Micu1*$^{ECKO}$ mice and *Micu1*$^{Fl/Fl}$ mice were shown in Figure S5.

#### Generation of endothelial cells-specific *Micu1* transgenic mice

Rosa26-CAG-LSL-Micu1-WPRE-polyA (*Rosa26*$^{LSL\text{-}Micu1}$) mice were generated on C57BL/6J background by Shanghai Model Organisms Center (Shanghai, China). CRISPR/Cas9 technology was used to obtain *Micu1* conditional expression mouse model of Rosa26 site-specific knock-in at chromosome 6 in C57BL/6J background. Guide RNA, Cas9 mRNA, targeting vector to recombination were injected to fertilized eggs by microinjection to obtain Rosa26-CAG-LSL-Micu1-WPRE-polyA

(*Rosa26*$^{LSL\text{-}Micu1}$) mice. Endothelial cells-specific *Micu1* transgenic mice (*Micu1*$^{ECTg}$) were obtained by crossbreeding *Rosa26*$^{LSL\text{-}Micu1}$ mice with *Cdh5*-Cre mice. *Rosa26*$^{LSL\text{-}Micu1}$ mice labeled as *Micu1*$^{WT}$ in this article were used for controls. The identification of *Micu1*$^{ECTg}$ mice and *Micu1*$^{WT}$ mice were shown in Figure S6.

**Human tissue samples**

The collection of human aortic samples or serum were approved by the institutional review board (IRB) of The First Affiliated Hospital of University of Science and Technology (No. 2021 KY 089). All experiments were performed in accordance with the relevant guidelines and regulations. Human tissues and serum were all collected from The First Affiliated Hospital of University of Science and Technology of China. Human aortic samples were obtained from patients with coronary artery disease who underwent aortic surgery (n=4). Serum samples of healthy people were from physical examination center of The First Affiliated Hospital of University of Science and Technology of China, and serum samples of people with coronary artery disease were collected from vascular surgery. The detailed information of human serum was presented in Table S1.

**Cell culture and treatments**

Different donors of human umbilical vein endothelial cells (HUVECs) were purchased from Lifeline Cell Technology (San Diego, CA, Cat# FC-0003, Lot 08119/10128/04608). HUVECs were cultured in Endothelial Cell Medium (ScienCell, Cat# 1001) consists of 500 ml of basal medium, 25 ml of fetal bovine serum (ScienCell, Cat# 0025), 5 ml of endothelial cell growth supplement (ScienCell, Cat# 1052) and 5 ml of antibiotic solution (penicillin /streptomycin, ScienCell, Cat# 0503).

Different donors of human aortic endothelial cells (HAECs) were from Lifeline Cell Technology (Cat# FC-0014, Lot 02357/01921/04675) and cultured in endothelial cell culture medium (Lifeline Cell Technology, Cat# LL-0013).

HUVECs or HAECs were 80% confluent treatment with different stimuli including lipopolysaccharides (Sigma-Aldrich, Cat# L2630), recombinant human TNFα (PeproTech, Cat# 300-01A), recombinant human IL-1β (PeproTech, Cat# 200-01B) for indicated time points.

**Transcriptomic profiling**

HUVECs were transfected with negative control siRNA or MICU1 siRNA (40 nM, RiboBio, Guangzhou, China) for 48 h. The sense sequence of MICU1 siRNA was (5'-3'): GCAGCUCAAGAAGCACUUCAA. Trizol (Invitrogen, Cat# 15596026) was used to extract the cells in control group and the treatment group. The strand-specific mRNA library building kit was used to generate mRNA libraries at high-throughput sequencing on the BGISEQ-500 (BGI) platform (Gene Denovo, Guangzhou, China). The purified sequences were compared with the human genome from NCBI by hierarchical indexing for spliced alignment of transcripts (HISAT). Based on the pure sequences that met the quality control standards, the differentially expressed genes were screened according to the fold change (FC) ≥2 and false discovery rate (FDR) <0.05. The obtained differences were compared with the Gene Ontology (GO) database and Kyoto Encyclopedia of Genes and Genomes (KEGG) database to enrich gene functional attributes and pathways to clarify the effect of MICU1 downregulation on the overall biological function of endothelial cells. The raw sequence data have been deposited in the Genome Sequence Archive (Genomics, Proteomics & Bioinformatics 2021) in National Genomics Data Center (Nucleic Acids Res 2022), China National Center for Bioinformation / Beijing Institute of Genomics, Chinese Academy of Sciences (GSA-Human: HRA005700, https://ngdc.cncb.ac.cn/gsa-human).

**RNA extraction and quantitative real-time-PCR**

Total RNA was extracted from endothelial cells using Trizol (Invitrogen, Cat. No.15596026). The collected RNA was converted to cDNA using a reverse transcription kit (TaKaRa, Japan, Cat# RR036A-1). quantitative realtime-PCR (qRT-

PCR) was performed with PCR System of LightCycler 96 (Roche) using TB Green Premix (TaKaRa, Japan, Cat# RR820) for amplification reaction. Relative expression of target gene was normalized to gene GAPDH. The sequences of primers used were listed in Table S3.

**RT-PCR**

Regular RT-PCR was performed using 2 × Taq Master Mix (Vazyme, China, Cat# P112-01) to identify the genotype of genetically modified mice. Amplification products amplified by ProFlex PCR system (Thermo Fisher Scientific) were separated in 1% agarose gel and analyzed by Gel Imaging System of JS-680D (P&Q Science Technology, China). The primer sequence used is as below:

*Micu1*$^{Fl/+}$ primer sequence (5'-3'): P1, ACCCGACTAAAGAGCAGCTT (forward); P2, CCAGCTCAGAGGAGCACTAA (reverse).

*Rosa26*$^{LSL\text{-}Micu1}$ primer sequence (5'-3'): P1, TCAGATTCTTTTATAGGGGACACA (forward); P2, TAAAGGCCACTCAATGCTCACTAA (reverse); P3, GTTGCGTCAGCAAACACAGT (forward); P4, ACTGTGATGGCAATGGGGAG (reverse).

*Cdh5*-Cre primer sequence (5'-3'): P1, CCAGGCTGACCAAGCTGAG (forward); P2, CCTGGCGATCCCTGAACA (reverse); P3, AGTGGCCTCTTCCAGAAATG (forward); P4, TGC GACTGTGTCTGATTTCC (reverse).

**Western blot**

Total protein was extracted using lysis buffer (Beyotime, China, Cat# P0013). The protein samples containing loading buffer (Beyotime, Cat# P0015) was subjected to SDS-PAGE electrophoresis. Then the proteins were transferred to PVDF membrane (Millipore, Cat# IPVH00010). The blocking buffer (LI-COR, Cat# 927-60001) was used to block, and the primary antibody was incubated overnight at 4℃. The information of primary antibody in detail was shown in Table S2. Next day, the fluorescent secondary antibody concerning IRDye® 800CW Goat anti-Rabbit IgG

(LI-COR, Cat# 925-32211) or IRDye® 680RD Donkey anti-Mouse IgG (LI-COR, Cat# 926-68072) was incubated. Images were detected by the Odyssey Infrared Imaging System (LI-COR) and analyzed by Image J software.

#### *En face* immunofluorescence staining

Mice were anesthetized by continuous inhalation of 2.5% isoflurane gas and fixed with normal saline containing 40 U/ml heparin followed a slow drip of precooled 4% paraformaldehyde (PFA) solution for 15 min. The aorta was isolated with cleaned of peri-adventitial tissue and cut longitudinally, which then penetrated in 0.1% Triton X-100 for 10 min, and blocked in 10% goat serum containing 2.5% Tween-20 for 1 h. The primary antibodies including MICU1 antibody (Sigma-Aldrich, Cat# HPA037480) and VE-cadherin (BD Biosciences, Cat# 555289) were then incubated overnight. Samples were incubated with Alexa Fluor™ 488-conjugated secondary antibodies (Invitrogen, Cat# A11001) or Alexa Fluor™ 546-conjugated secondary antibodies (Invitrogen, Cat# A11010), and the nuclei were stained with DAPI (Beyotime, Cat# C1006) for 5 min. The images were detected by Leica STED high-resolution laser confocal system (Leica, Germany).

#### Evaluation of atherosclerosis

Male and female *Micu1*$^{ECKO}$ mice, male *Micu1*$^{ECTg}$ mice and littermate control mice about 8-week-old were administered a tail vein injection of adeno-associated virus expressing mouse recombinant PCSK9 with mutation of D377Y (AAV8-PCSK9$^{D377Y}$, Vigene Biosciences, China) with $5\times10^{11}$ viral particles in a 100 μL volume of sterile saline. The methods for induction of atherosclerosis were first described by Martin Maeng Bjørklund et. al (Circ Res. 2014 May 23;114(11):1684-9). Then mice were fed with western diet (Research Diets, Cat# D12108C) for 12 weeks. After that, mice were anesthetized by continuous inhalation of 2.5% isoflurane gas. Hearts and whole aortas were isolated and then fixed in 4% PFA solution. The whole aortas were subjected Oil Red O (Poly Scientific R&D, Cat# s1849-16OZ) staining to quantification of atherosclerotic lesion which calculated by Image J software. The aortic root sections were made to value aortic sinus plaque. The sections were stained with Oil Red O solution and snapped by ZEISS Axio Imager (ZEISS, Germany). The positive area of Oil Red O staining was quantified by Image J software.

**Measurements of lipid profile in mouse serum**

Blood was collected retroorbitally from the mice after overnight fasting. Then the blood was centrifuged to obtain serum. Serum triglyceride (TG), cholesterol (CHO), high-density lipoprotein (HDL) and low-density lipoprotein (LDL) were detected using automatic biochemical analyzer of Chemray (Rayto, Shenzhen, China).

**Immunofluorescence staining and confocal microscopy**

For immunofluorescence staining of sections of thoracic aorta or aortic sinus, the sections were permeabilized with 0.1% Triton X-100 at room temperature for approximately 15 min. Then the sections were blocked with the immunostaining blocking solution (Beyotime, China, Cat# P0102) at room temperature for 1 h. After the blocking solution was discarded, the primary antibodies of ICAM1 (Abcam, Cat# ab222736), VCAM1 (Abcam, Cat# ab134047), SIRT3 (Cell Signaling Technology, Cat# 5490), SOD2 (Cell Signaling Technology, Cat# 5490) and Ac-SOD2 (Abcam, Cat# ab137037) were added and placed at 4 °C overnight. The following steps were performed in the dark. The secondary fluorescent antibodies were added and incubated for 1 h, and then nuclei were stained with DAPI staining solution (Beyotime, Cat# C1006) for 5 min. Images were examined using Leica STED high-resolution laser confocal system (Leica, Germany).

For immunofluorescence staining of cells, cells were seeded in confocal dishes for imaging. Mitochondria were stained with Mito-Tracker Red CMXRos (Beyotime, Cat# C1035). Then cells were fixed with 4% PFA solution and permeabilized with 0.1% Triton X-100 at room temperature for approximately 5 min. The following steps were similar to those above.

#### Histopathology

Frozen cross-sections from fixed aortic sinus were stained with hematoxylin and eosin (H&E, Servicebio, China, Cat# G1076), Masson's trichrome (Servicebio, China, Cat# G1006) according to the manufacturer's protocols. Images were captured by ZEISS Axio Imager (ZEISS, Germany).

#### Enzyme-linked immunosorbent assay

The cytokines were tested using enzyme-linked immunosorbent assay (ELISA) kits. Blood was collected from the retroorbital of the mice. Then the blood was centrifuged to obtain serum. ELISA analysis detected serum IL-6 (Boster, China, Cat# EK0411), serum TNFα (Boster, Cat# EK0527), serum MCP-1 (Boster, Cat# EK0568), serum CXCL-10 (Boster, Cat# EK0736) and serum E-selectin (Boster, Cat# EK0502) according to the manufacturer's introductions. The results were obtained using multifunctional microplate reader of SpectraMax iD3 (Molecular Devices, USA).

#### siRNA transfection and adenovirus infection

Cells were planted in dishes at a density of about $5\times10^5$/ml and then allowed to grow to about 60% cell density for experiments. Hiperfect transfection reagent (QIAGEN, Cat# 301705) was used to prepare small interfering RNA (RiboBio, Guangzhou, China) system with final concentration of 20 nM, 40 nM or 80 nM. The Hiperfect transfection mixture was added to the dish in Opti-MEM medium (Gibco, Cat# 31985088), cultured for 6 h, then the solution was aspirated, and ECM medium was added to culture for 48 h. The efficiency of knockdown was detected by western blot.

Cells were seeded in dishes, and experiments were performed when the cell density had grown to about 80%. The medium in the dish was replaced by Opti-MEM containing 1% fetal bovine serum (FBS), and an amount of MICU1 adenovirus (Vigene Biosciences, China) with final concentration of 50 MOI, 100 MOI or 200 MOI was added to it. After 4-6 h of culture, the solution was changed to ECM medium for 48 h. MICU1 overexpression was detected by western blot.

**Measurement of mitochondria $Ca^{2+}$**

Rhod 2-AM (Invitrogen, Cat# R1245MP) mitochondrial $Ca^{2+}$ fluorescent probe was used to measure mitochondria $Ca^{2+}$. Pluronic F127 was added to Rhod 2-AM solution to prevent Rhod 2-AM from polymerizing in HBSS solution and facilitated its entry into cells. Rhod 2-AM was diluted with HBSS and added to the cells at a final concentration of 5 μM. The cells were incubated at 37℃ for 20 min, then HBSS containing 1%FBS was added for 40min. $[Ca^{2+}]_m$ was observed using a ZEISS laser confocal microscope LSM980 (ZEISS, Germany). After 1 min of baseline recording, histamine (HT, 50 μM, MedChemExpress, Cat# HY-B1204) was added. The fluorescence intensity was detected in real time every 600 ms for 300 s at 561nm excitation.

**Detection of mitochondria ROS**

For confocal microscopy, MitoSox (Invitrogen, Cat# M36008) fluorescent probe was used for detection of reactive oxygen species (ROS). The working solution diluted in HBSS with a final concentration of 5 μM, added to the test cells, and incubated at 37 ℃ in the dark for 10 min. The fluorescence intensity was detected by ZEISS laser confocal microscope LSM980 (ZEISS, Germany) at 510/580 nm. For flow cytometry, after MitoSox incubation, cells were digested and placed in saline and examined by flow cytometry CytoFLEX (Beckman, USA).

**Measurement of ROS**

Dihydroethidium (DHE, Beyotime, Cat# S0063) fluorescent probe was used for detection. The working solution of DHE was diluted to a final concentration of 1 μM, which then was added to the cells and incubated at 37 °C in the dark for 30 min. The fluorescence intensity was detected by ZEISS laser confocal microscope LSM980 (ZEISS, Germany) at 510/580 nm.

**Measurement of NO**

DAF-FM DA means 3-Amino,4-aminomethyl-2',7'-difluorescein, diacetate (Beyotime, Cat# S0019) used to detect the level of nitric oxide (NO). DAF-FM DA diluted to a final concentration of 5 μM was added to the cells and then incubated at 37 °C in the dark for 30 min. The fluorescence intensity was detected by ZEISS laser confocal microscope LSM980 (ZEISS, Germany) at 488/507 nm.

**SOD2 activity**

SOD2 activity was measured using the CuZn/Mn-SOD activity Detection kit (Beyotime, Cat# S0103) related to WST-8 method. The detection was performed according to the manufacturer's protocols. Briefly, cells were collected and homogenized in cold phosphate buffered saline (PBS), then centrifuged to obtain the supernatant as the sample to be tested. Samples and various blank controls were added into 96 well plate, then added SOD detection buffer, WST-8 enzyme working solution, and reaction start working solution in sequence, which were incubated at 37 °C for 30 min before detecting absorbance at 450 nm using multifunctional microplate reader of SpectraMax iD3 (Molecular Devices, USA).

#### Supplemental Tables

**Table S1 Characteristics of CAD patients and controls included in this study.**

| Parameter | CAD (n=13) | Controls (n=11) | p Value |
|---|---|---|---|
| Male/female | 12/1 | 8/3 | 0.3002 |
| Age, y | 66.9±11.7 | 40.2±15.8 | <0.0001 |
| CAD, n (%) | 13 (100) | 0 (0) | N/A |
| Hypertension, n (%) | 10 (76.9) | 1 (9.1) | 0.0013 |
| Diabetes, n (%) | 7 (53.8) | 0 (0) | 0.0059 |
| TC (mmol/L) | 3.87±0.90 | 5.27±0.85 | 0.0008 |
| TG (mmol/L) | 1.32±0.82 | 2.54±2.99 | 0.1711 |
| HDL (mmol/L) | 0.99±0.36 | 1.41±0.28 | 0.0047 |
| LDL (mmol/L) | 2.38±0.70 | 3.16±0.51 | 0.0056 |

CAD, coronary artery disease; N/A, not applicable; TC, total cholesterol; TG, triglyceride; HDL, high-density lipoprotein; LDL, low-density lipoprotein.

**Table S2 List of antibodies for Immunoblotting and immunofluorescence.**

| **Antibodies for western blot** | | | | |
|---|---|---|---|---|
| **Name** | **Source** | **Cat. No.** | **Species** | **Titer** |
| MICU1 | Sigma-Aldrich | HPA037480 | Rabbit | 1:500 |
| MCU | Cell Signaling Technology | 14997 | Rabbit | 1:1000 |
| VDAC1 | Santa Cruz Biotechnology | sc-390996 | Mouse | 1:250 |
| β-actin | Proteintech | 66009-1-Ig | Mouse | 1:5000 |
| SIRT3 | Cell Signaling Technology | 5490 | Rabbit | 1:1000 |
| VCAM1 | Abcam | ab134047 | Rabbit | 1:1000 |
| Ac-SOD2 | Abcam | ab137037 | Rabbit | 1:800 |
| SOD2 | Cell Signaling Technology | 13141 | Rabbit | 1:1000 |
| α-Tubulin | Proteintech | 66031-1-Ig | Mouse | 1:5000 |

| | | | | |
|---|---|---|---|---|
| IRDye® Goat anti-Rabbit IgG | LI-COR | 925-32211 | Goat | 1:10000 |
| IRDye® Donkey anti-Mouse IgG | LI-COR | 926-68072 | Donkey | 1:10000 |
| **Antibodies for immunofluorescence** | | | | |
| Name | Source | Cat. No. | Species | Titer |
| MICU1 | Sigma-Aldrich | HPA037480 | Rabbit | 1:50 |
| VE-cadherin | BD Biosciences | 555289 | Rat | 1:100 |
| ICAM1 | Abcam | ab222736 | Rabbit | 1:100 |
| VCAM1 | Abcam | ab134047 | Rabbit | 1:100 |
| CD68 | Bio-Rad | MCA1957GA | Rat | 1:100 |
| α-SMA | Proteintech | 14395-1-AP | Rabbit | 1:100 |
| SIRT3 | Cell Signaling Technology | 5490 | Rabbit | 1:100 |
| Ac-SOD2 | Abcam | ab137037 | Rabbit | 1:50 |
| SOD2 | Proteintech | 24127-1-AP | Rabbit | 1:100 |
| Alexa Fluor™ 488 Goat anti-Rabbit IgG | Beyotime | A0423 | Goat | 1:1000 |
| Alexa Fluor™ 488 Goat anti-Mouse IgG | Invitrogen | A-11001 | Goat | 1:1000 |
| Alexa Fluor™ 546 Goat anti-Rabbit IgG | Invitrogen | A-11010 | Goat | 1:1000 |
| Alexa Fluor™ 488 Goat anti-Rat IgG | Invitrogen | A-11006 | Goat | 1:1000 |

**Table S3 List of primers for qRT-PCR.**

| Description | Species | Sequence (5'-3') |
|---|---|---|
| MICU1 | Mouse | forward: CTTGAATGGAGACGGAGAGG<br>reverse: ACTTGAGGGTGTTCCCAGTG |
| GAPDH | Mouse | forward: GGTTGTCTCCTGCGACTTCA<br>reverse: TGGTCCAGGGTTTCTTACTCC |
| MICU1 | Human | forward: GTGTTCAGCCCTCACAACCT |

| | | |
|---|---|---|
| | | reverse: TCTCCCATCCACAGGGTCAT |
| CXCL1 | Human | forward:CCCAAGAACATCCAAAGTGTGA<br>reverse: CAAGCTTTCCGCCCATTCT |
| CXCL2 | Human | forward: CCTGCAGGGAATTCACCTCA<br>reverse: TGAGACAAGCTTTCTGCCCA |
| CXCL3 | Human | forward: GTGAATGTAAGGTCCCCCGG<br>reverse: TTCTGAACCATGGGGGATGC |
| CX3CL1 | Human | forward: GGGCGTCCTTATCACTCCTG<br>reverse: TGGTAGGTGAACATGGCCAC |
| IL12A | Human | forward: AGTTTGGCCAGAAACCTCCC<br>reverse: TTTGTCTGGCCTTCTGGAGC |
| IL15 | Human | forward: GTTTCAGTGCAGGGCTTCCT<br>reverse: TGCAACTGGGGTGAACATCA |
| CCL5 | Human | forward: GTGTGCCAACCCAGAGAAGA<br>reverse: GCCTCCCAAGCTAGGACAAG |
| VEGF-C | Human | forward: GCAGTTACGGTCTGTGTCCA<br>reverse: TCCTTGAGTTGAGGTTGGCC |
| MYD88 | Human | forward: ACTTGGAGATCCGGCAACTG<br>reverse: TTGGTAAGCAGCTCGAGCAG |
| CSF1 | Human | forward: CCAGCAACTTCCTCTCAGCA<br>reverse: GGGGTGTGATTCCAGTCCTG |
| IFNβ1 | Human | forward: GCCGCATTGACCATCTATGA<br>reverse: GCCAGGAGGTTCTCAACAATAG |
| IL6 | Human | forward: TGCAATAACCACCCCTGACC<br>reverse: AGCTGCGCAGAATGAGATGA |
| CXCL10 | Human | forward: TGTACGCTGTACCTGCATCA<br>reverse: GCAATGATCTCAACACGTGGA |
| VCAM1 | Human | forward: TGTCAATGTTGCCCCCAGA<br>reverse: TGCTCCACAGGATTTTCGGA |
| TNFα | Human | forward: GCACTGAAAGCATGATCCGG<br>reverse: AGAGGCTGAGGAACAAGCAC |
| MCP1 | Human | forward: TGTCCCAAAGAAGCTGTGATC<br>reverse: ATTCTTGGGTTGTGGAGTGAG |

| | | |
|---|---|---|
| GAPDH | Human | forward: CACCATCTTCCAGGAGCGAG<br>reverse: CCTTCTCCATGGTGGTGAAGAC |
| MICU2 | Human | forward: GGCAGTTTTACAGTCTCCGC<br>reverse: AAGAGGAAGTCTCGTGGTGTC |
| MICU3 | Human | forward: TCGCAGAAACACCACCAGTT<br>reverse: GAACCCTGCATGTGGCTTTG |
| MCU | Human | forward: CCTCCCAAAGTGCCGAGATT<br>reverse: ACCCCATAGCACCAAAGTGG |
| MCUb | Human | forward: CCCAGGTTTTGCGTGTGAAG<br>reverse: GTGTTACCAAGGGAAGGCCA |
| EMRE | Human | forward: CTCGCTGGCTAGTATTGGCA<br>reverse: AACGATGACTGACCTCGACG |

### Supplemental Figures

Figure S1

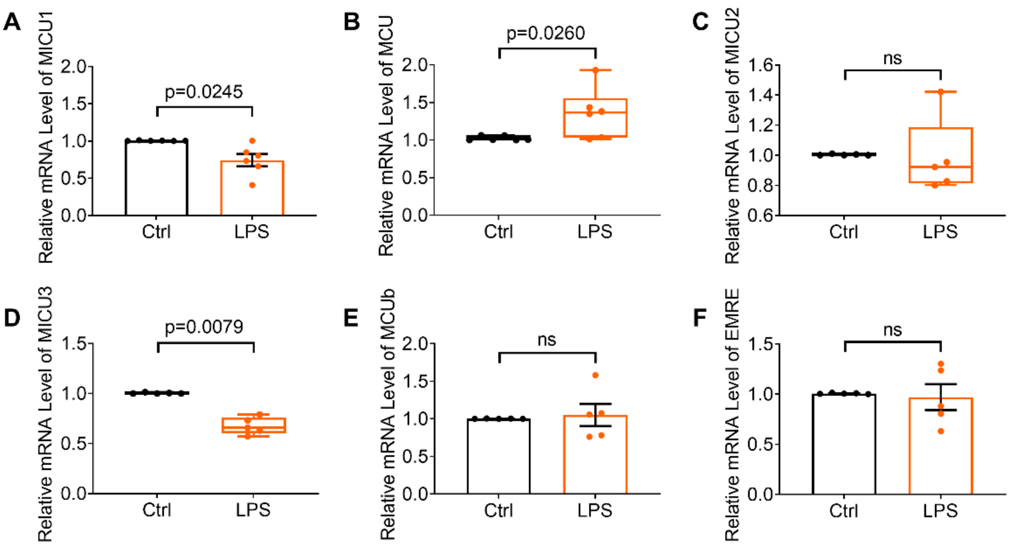


**Figure S1. The mRNA level of mitochondrial $Ca^{2+}$ uniporter complex in response to LPS treatment.**

mRNA level of MICU1 (A), MCU (B), MICU2 (C), MICU3 (D), MCUb (E), EMRE (F) in response to LPS treatment in HUVECs.

Statistical analysis was performed by Welch's *t* test (A, E, F) and Mann-Whitney *U* test (B-D).

Figure S2

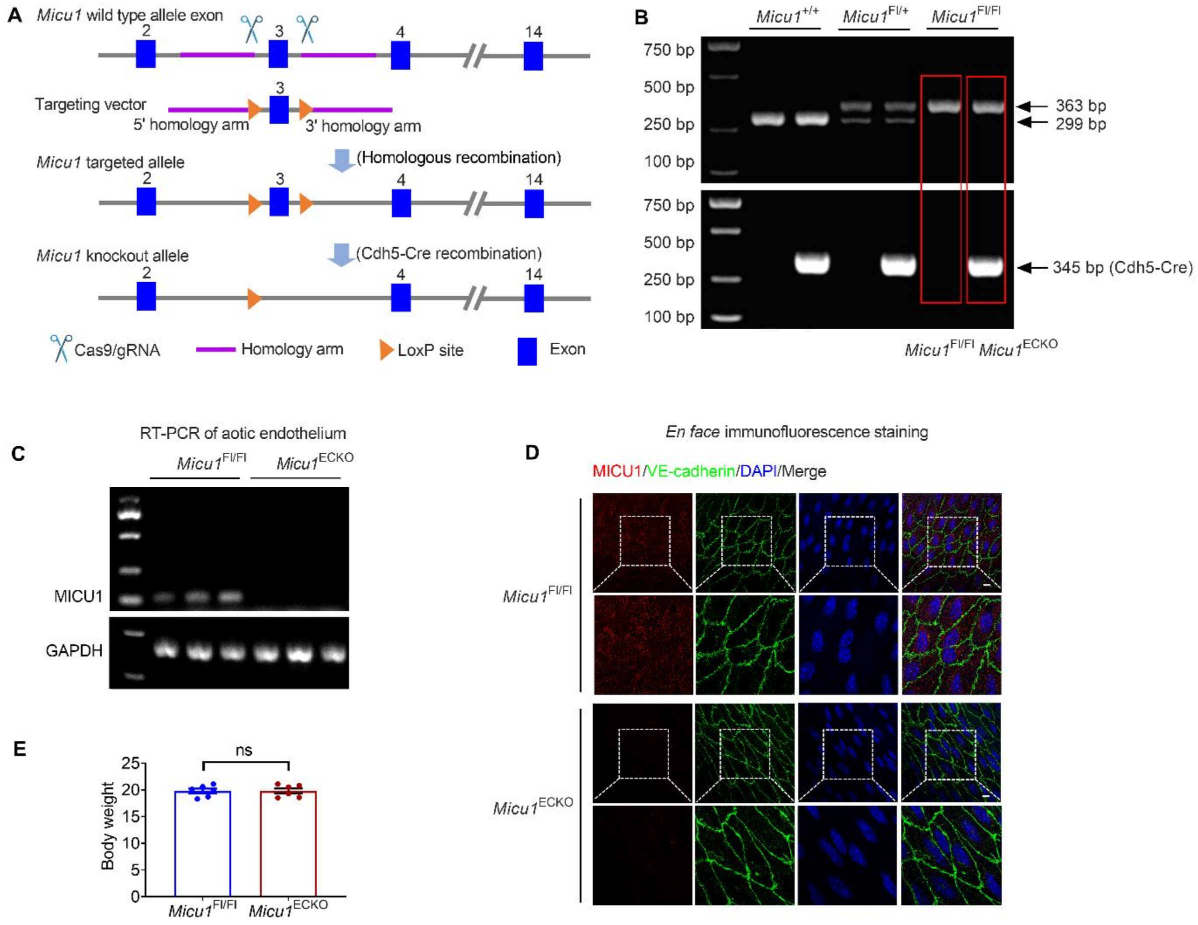


**Figure S2. Generation and validation of endothelial-specific *Micu1* knockout mice.**

A. Breeding strategy of endothelial-specific *Micu1* knockout (*Micu1*$^{\mathrm{ECKO}}$) mice.

B. Mouse genomic DNA were isolated for genotyping *Micu1*$^{\mathrm{Fl/Fl}}$ mice and *Micu1*$^{\mathrm{ECKO}}$ mice.

C. The mRNA level of MICU1 was measured by RT-PCR in mouse aortic endothelial cells.

D. *En face* immunofluorescence staining showed the protein expression of MICU1 was ablated in *Micu1*$^{\mathrm{ECKO}}$ mouse aorta (n=5). Scale bars, 10 μm.

E. Body weight of *Micu1*$^{\mathrm{Fl/Fl}}$ mice and *Micu1*$^{\mathrm{ECKO}}$ mice (n=6). Statistical analysis was performed by Student *t* test.

Figure S3

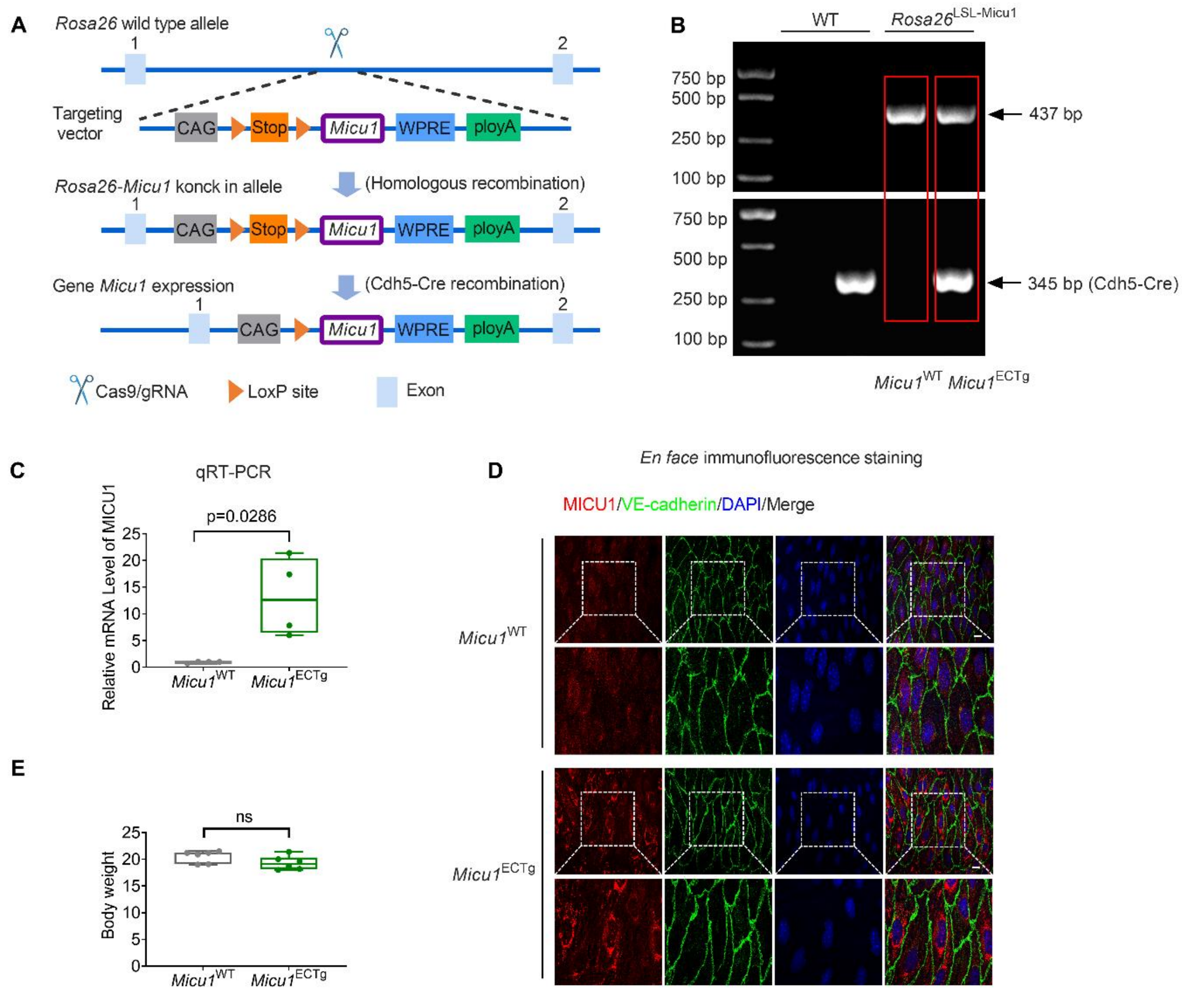


**Figure S3. Generation and validation of endothelial-specific *Micu1* trangenic mice.**

A. Breeding strategy of endothelial-specific *Micu1* trangenic (*Micu1*$^{ECTg}$) mice.

B. Mouse genomic DNA were isolated for genotyping *Micu1*$^{WT}$ mice and *Micu1*$^{ECTg}$ mice.

C. The mRNA level of MICU1 was measured by quantitative reverse transcription polymerase chain reaction (qRT-PCR) in mouse aortic endothelial cells (n=4).

D. *En face* immunofluorescence staining showed successful overexpression of MICU1 protein in *Micu1*$^{ECTg}$ mouse aorta (n=5). Scale bars, 10 μm.

E. Body weight of *Micu1*$^{WT}$ mice and *Micu1*$^{ECTg}$ mice (n=6).

Statistical analysis was performed by Student *t* test (C) and Mann-Whitney *U* test (F).

Figure S4

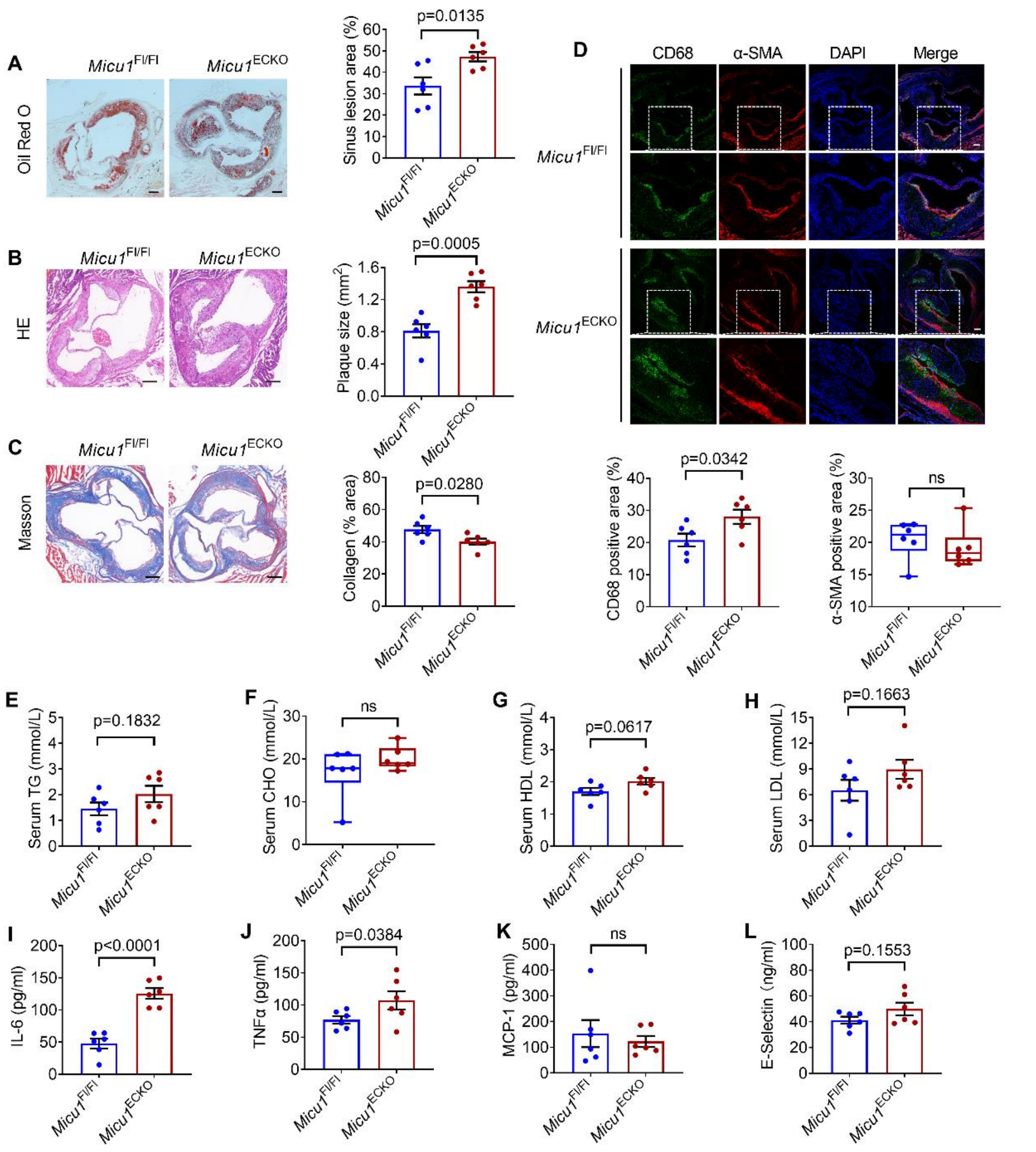


**Figure S4. *Micu1* deficiency in endothelial cells promoting the development of Western diet-induced atherosclerosis in female mice.**

A. Representative images of Oil Red O staining of atherosclerotic lesions of aorta in female *Micu1*$^{Fl/Fl}$ mice or *Micu1*$^{ECKO}$ mice infected with AAV8-PCSK9$^{D377Y}$ after 12 weeks of western diet (n=11). Scale bars, 1 mm.

B and C. Oil Red O staining (B) or hematoxylin-eosin staining (H&E) staining (C) of lesions of the aortic root in female *Micu1*$^{Fl/Fl}$ mice or *Micu1*$^{ECKO}$ mice from (A) (n=6). Scale bars, 200 μm.

D. Staining of CD68-positive macrophages in lesion area of the aortic sinus from female *Micu1*$^{Fl/Fl}$ mice or *Micu1*$^{ECKO}$ mice from (A) (n=6). Scale bars, 100 μm.

E-J. Serum levels of alanine aminotransferase (ALT), aspartate aminotransferase (AST, n=6 vs. 5), triglyceride (TG), cholesterol (CHO), high-density lipoprotein (HDL) and low-density lipoprotein (LDL) were detected in female *Micu1*$^{Fl/Fl}$ mice or *Micu1*$^{ECKO}$ mice infected with AAV8-PCSK9$^{D377Y}$ after 12 weeks of western diet (n=6).

K-N. ELISA of serum IL-6 (K), TNFα (L), MCP-1 (M), E-selectin (N) from female *Micu1*$^{Fl/Fl}$ mice or *Micu1*$^{ECKO}$ mice infected with AAV8-PCSK9$^{D377Y}$ after 12 weeks of western diet (n=6).

Statistical analysis was performed by Student *t* test (A-C, D, left panel, E, G-L) and Mann-Whitney *U* test (D, right panel, F).

Figure S5

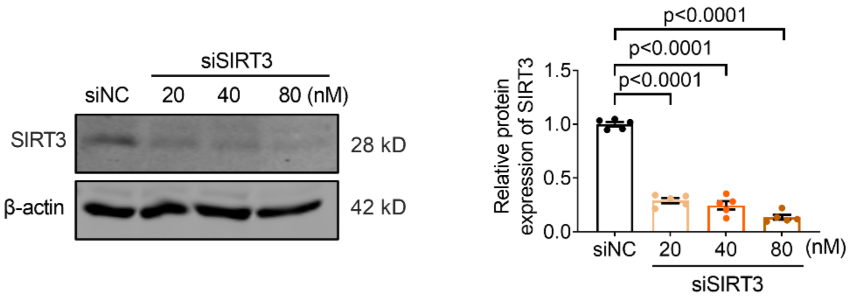


**Figure S5. The efficiency of SIRT3 knockdown.**

Expression of SIRT3 was detected by immunoblot in HUVECs treated with negative control siRNA (siNC) or SIRT3 siRNA (siSIRT3) for 48 h (n=5). Statistical analysis was performed by 1-way ANOVA followed by Bonferroni post hoc tests.
